## Supplementary material for "Production of germ-free mosquitoes via transient colonisation allows stage-specific investigation of host-microbiota interactions": Figure S

### Supplementary Information

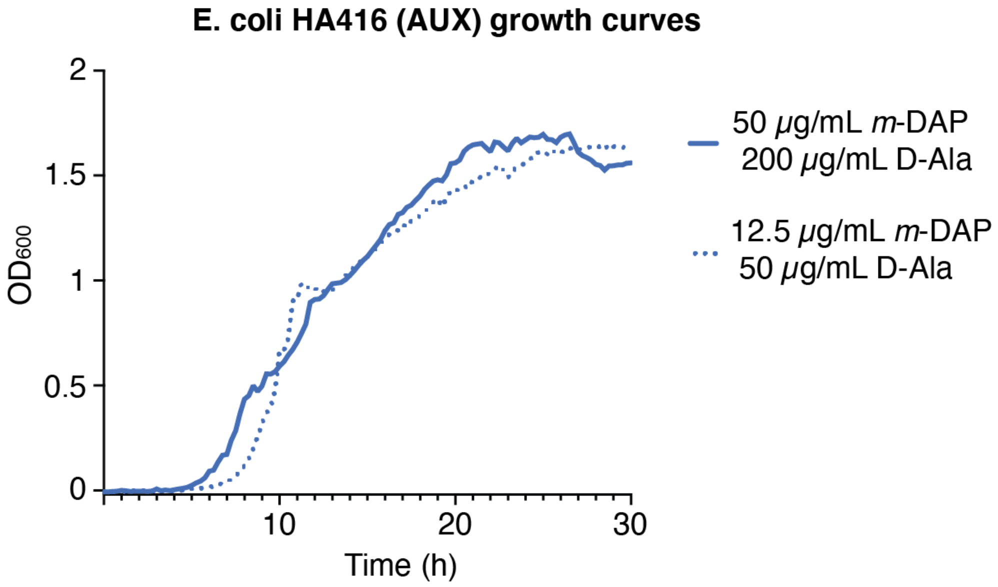


**Figure S1.** Growth curves of AUX *E. coli* in LB supplemented with optimal concentrations of *m*-DAP and D-Ala from [1] (continuous line) and concentrations compatible with larval growth (dotted line).

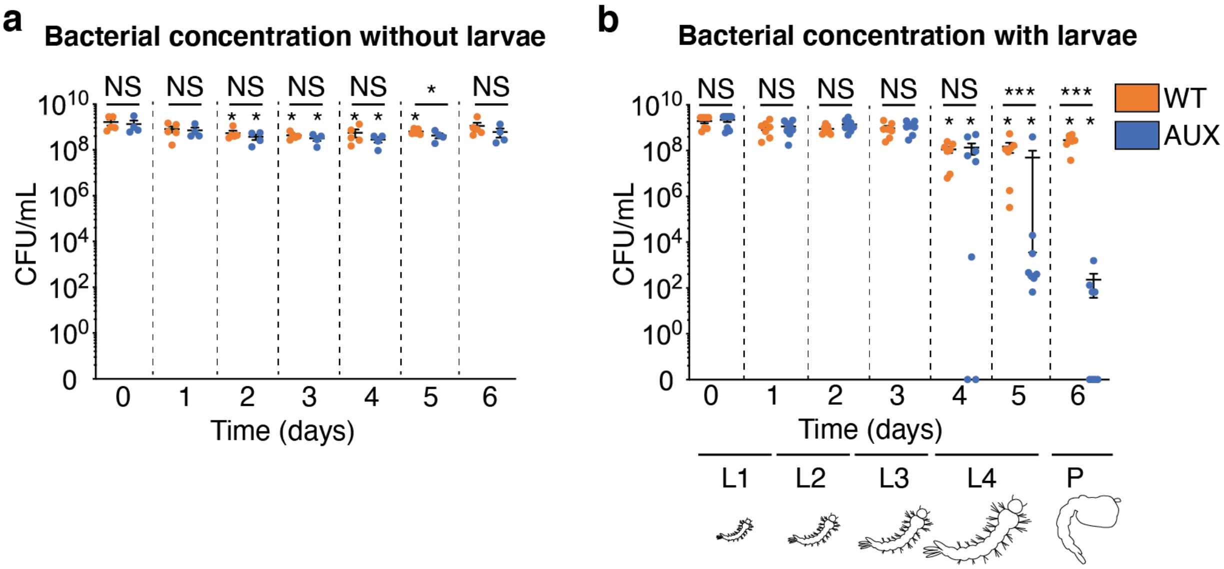


**Figure S2.** Bacterial concentration in larval rearing water in absence (**a**) and presence of larvae (**b**). Data show mean ± SEM of four independent replicates. Symbols above horizontal lines represent the level of significance of test comparing WT and AUX bacterial loads at each time-point. Asterisks below horizontal lines represent the level of significance comparing for each bacteria starting loads (T=0) with loads measured at each time point. Glmm with lsmeans (Bonferroni correction): NS, Non-significant; *, $p < 0.05$, **, $p < 0.01$, ***, $p < 0.001$.

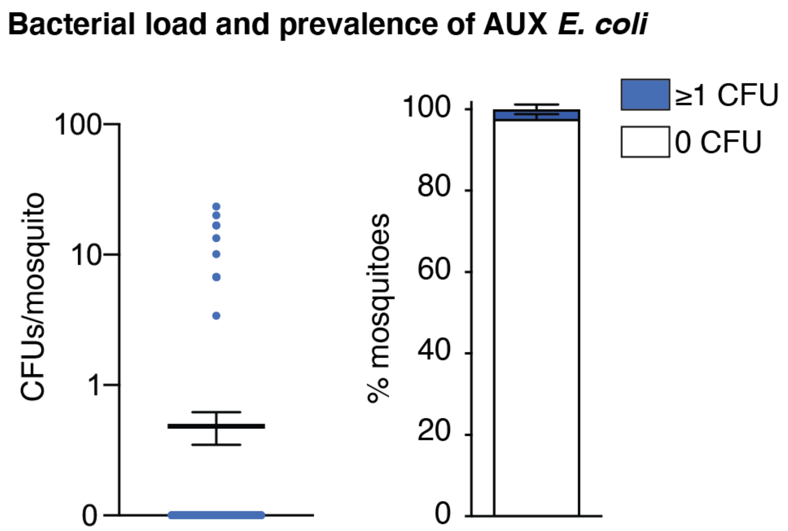


**Figure S3.** Bacterial load and prevalence of contaminated mosquitoes after reversible colonisation. Data represents mean ± SEM of three independent replicates where 34, 122 and 321 mosquitoes were analysed 0-2 days after emergence.

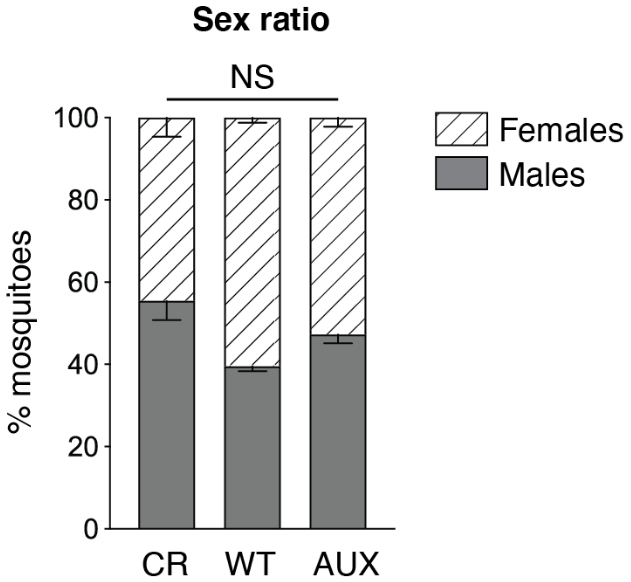


**Figure S4.** Sex ratio of conventionally reared and gnotobiotic mosquitoes. Ratio between adult females (striped bars) and males (full bars) emerged from conventionally-reared (CR), WT *E. coli* gnotobiotic (WT), and AUX *E. coli* gnotobiotic larvae (AUX, mean ± SEM). Data shown in Figure 1b and Figure S4 derive from the same experiments. Glmm: NS, non-significant.

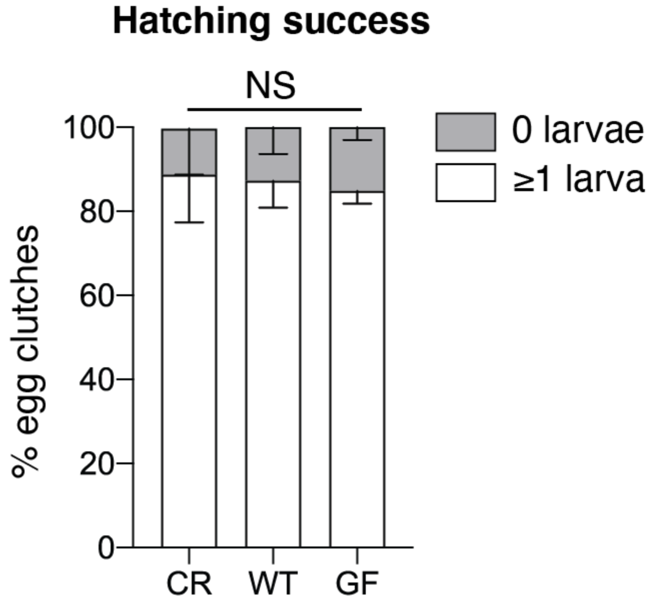


**Figure S5.** Percentage of egg clutches producing viable larvae (white bars) and not originating larvae (grey bars) for mosquitoes conventionally reared (CR), reared on WT *E. coli* (WT) and reversibly colonised (GF, mean ± SEM). Data shown in Figure 2d-e and Figure S5 derive from the same experiments. Glmm: NS, non-significant.

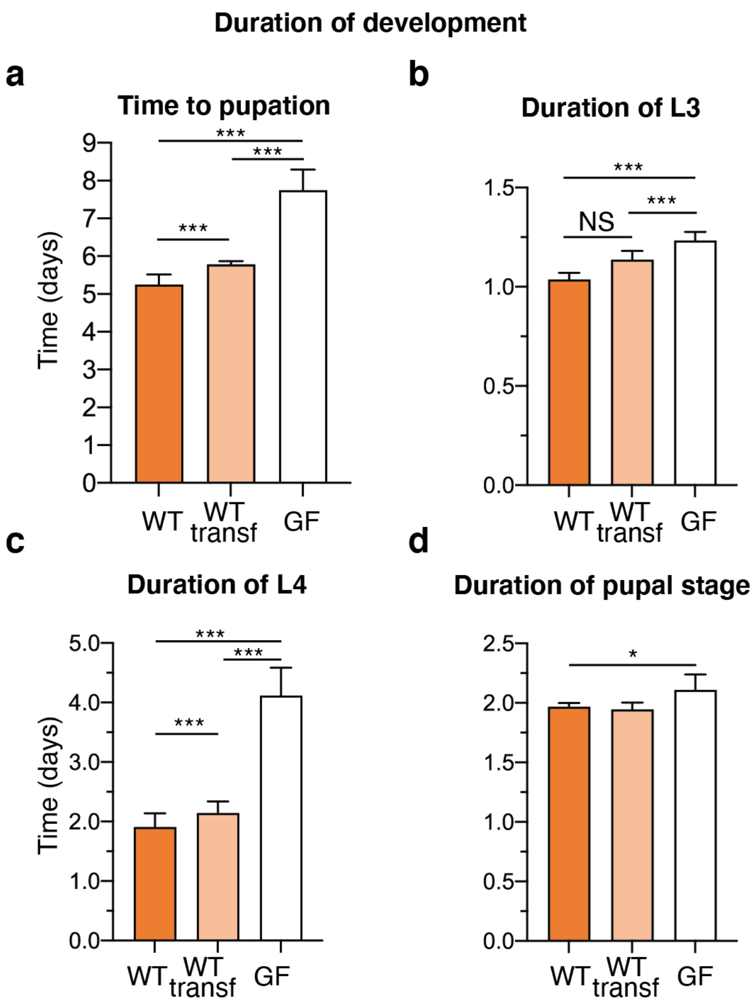


**Figure S6.** Duration of development of third instar larvae transferred in new sterile medium. Total time to pupation (**a**), duration of the third instar (L3, **b**), fourth instar (L4, **c**) and pupal stage (**d**) when continuously colonised (WT, orange), reared on WT *E. coli* and transferred in new rearing medium (WT transferred, light orange) and after becoming germ-free (GF, white, mean ± SEM). Data shown in Figure 3c and Figure S6 derive from the same experiments. Glmm with lsmeans (Bonferroni correction): NS, Non-significant; *, $p < 0.05$, **, $p < 0.01$, ***, $p < 0.001$.

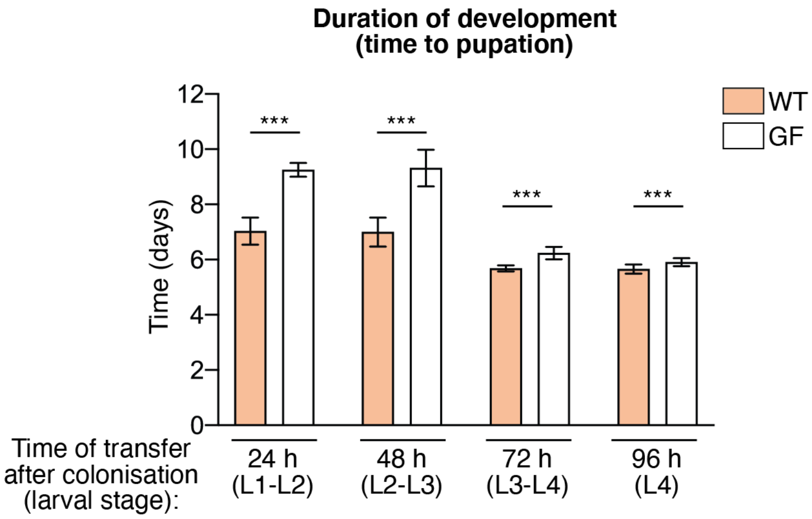


**Figure S7.** Duration of development of larvae transferred in new sterile medium at different time-points when reared on wild-type *E. coli* and transferred in new rearing medium (WT, light orange) and after becoming germ-free (GF, white, mean ± SEM). Data shown in Figure 3e and Figure S7 derive from the same experiments. Glmm: ***, $p < 0.001$.

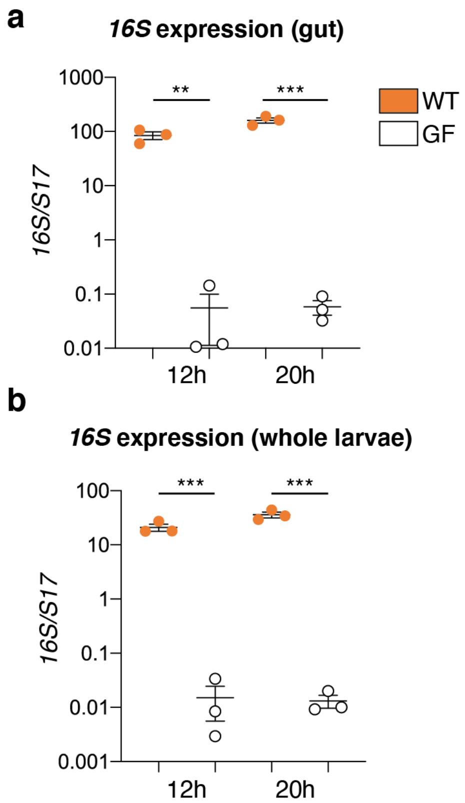


**Figure S8.** Quantification via qPCR of the bacterial load in guts (**a**) and whole larvae (**b**) of colonised and germ-free third-instar larvae at the 12 h and 20 h time-points. Data are expressed as the ratio between bacterial *16S* rRNA and *Ae. aegypti S17* genes in WT *E. coli*-carrying larvae (WT, orange) and in germ-free larvae (GF, white) and represents mean ± SEM of three independent replicates. The RNA samples were those analysed in the transcriptomic study (n = 60 larvae per condition per replicate). Glmm: **, $p < 0.01$; *** $p < 0.001$.

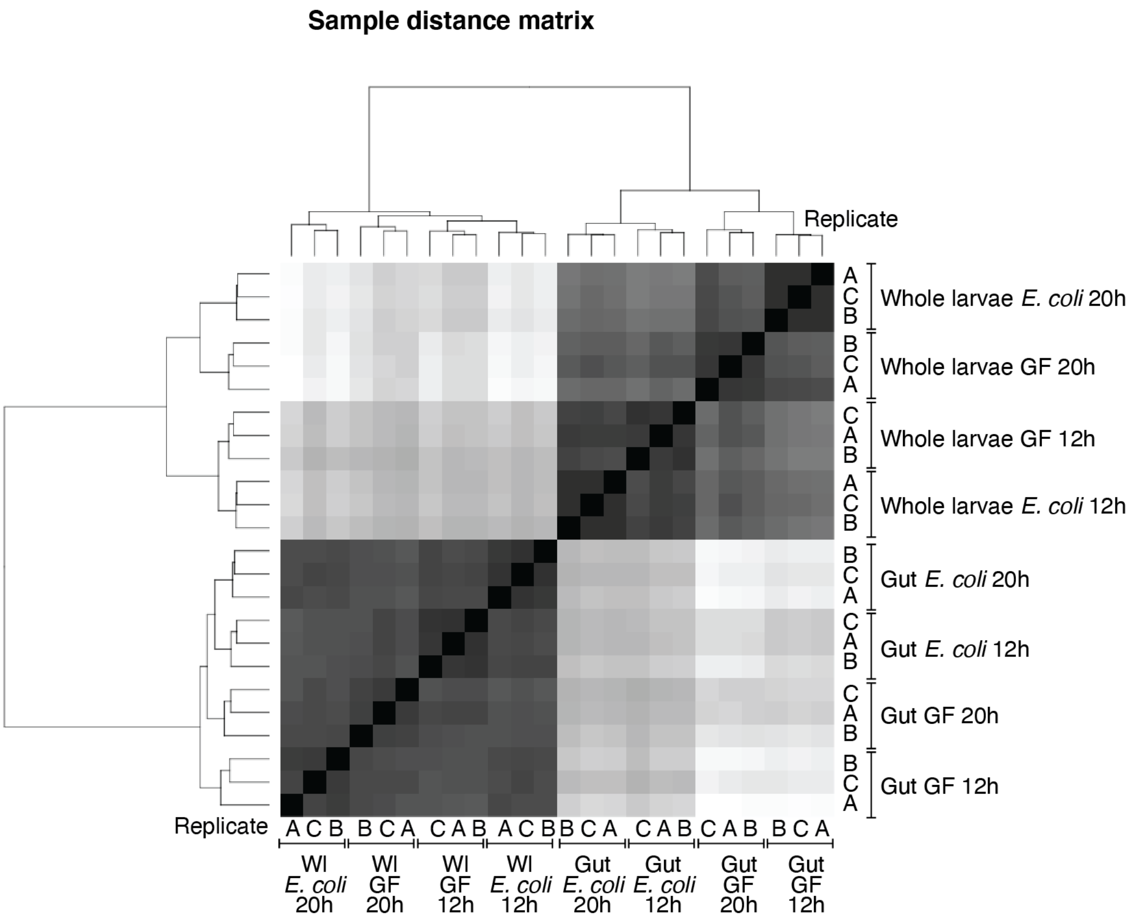


**Figure S9.** Sample distance matrix of gut and whole larvae transcriptomic data at the 12 h and 20 h time-points. Heatmap showing the regularized log transformation of the differential expression analysis data performed with DESeq2.

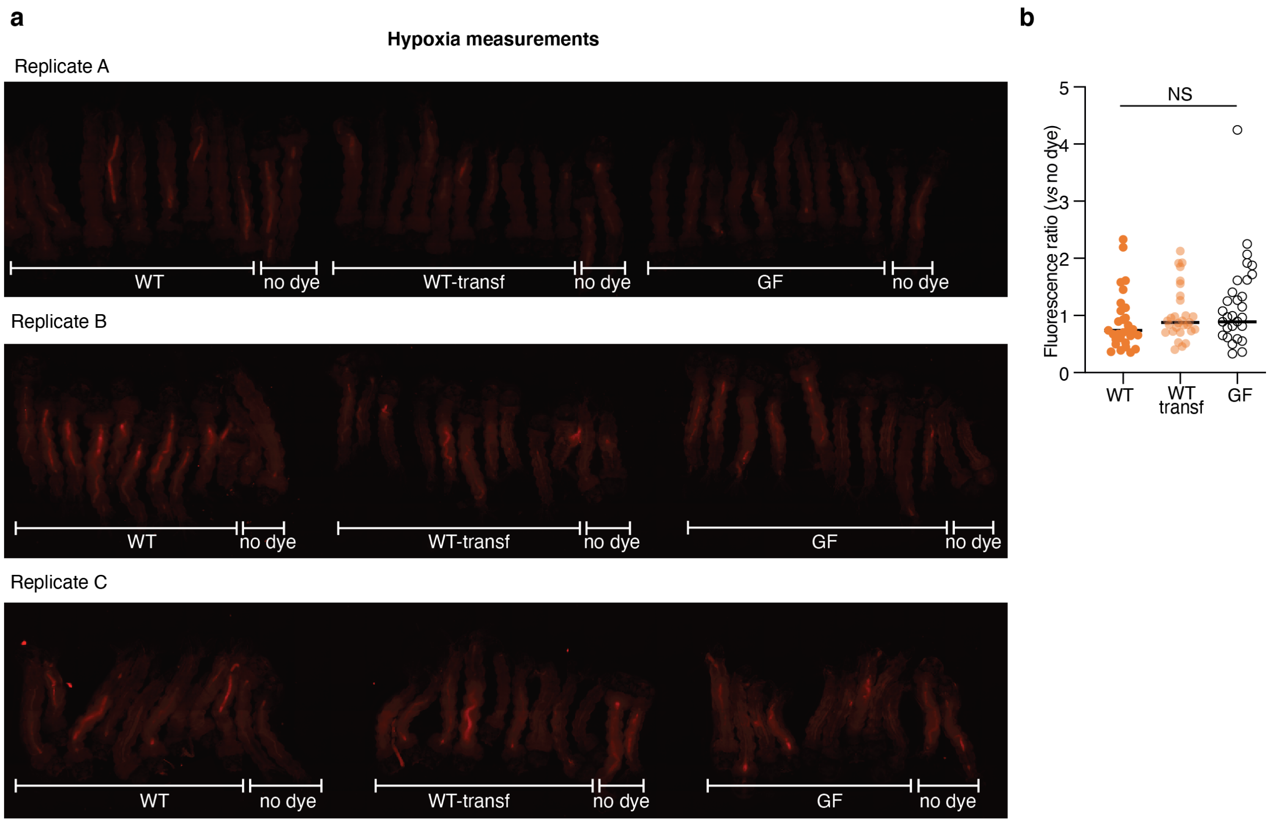


**Figure S10.** Hypoxia measurements on colonised and germ-free third-instar larvae 12 h after transfer. (**a**) Images show third-instar larvae gnotobiotic for *E. coli* WT before (WT) and after the transferring to new rearing medium (WT transf) and germ-free (GF) stained with the Image-iT Red Hypoxia Reagent at the 12 h time-point. For each condition, 9-10 larvae were stained with the dye, while 2 larvae were kept without dye as negative control for autofluorescence. For each larva, the ratio between the fluorescence intensity was normalised to the larval area. (**b**) Measurements of fluorescence intensity ratios with respect to no dye controls. Glmm: NS, Non-significant.

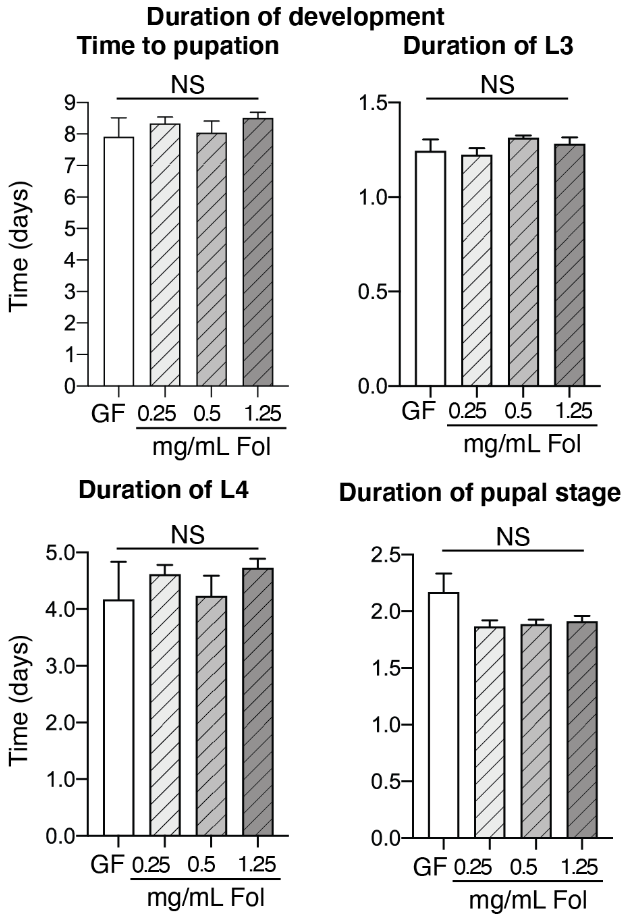


**Figure S11.** Duration of development of third-instar germ-free larvae with or without folate supplementation at different concentrations. Total time to pupation (**a**), duration of the third instar (L3, **b**), fourth instar (L4, **c**) and pupal stage (**d**, mean ± SEM). Data shown in Figure 6b and Figure S11 derive from the same experiments. Glmm: NS, Non-significant.

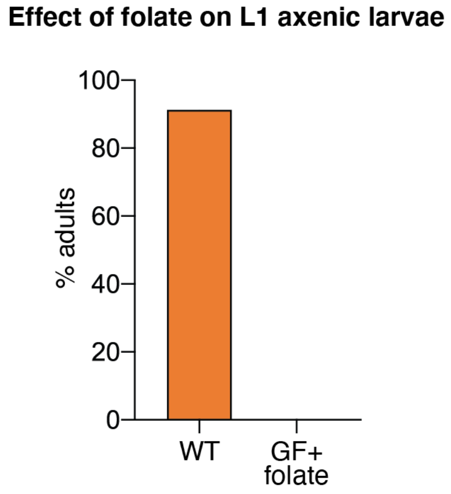


**Figure S12.** Effect of 1.25 mg/mL folate supplementation on the development of first instar axenic larvae (GF, n=24), compared to larvae colonised with WT *E. coli* (WT, n=48).

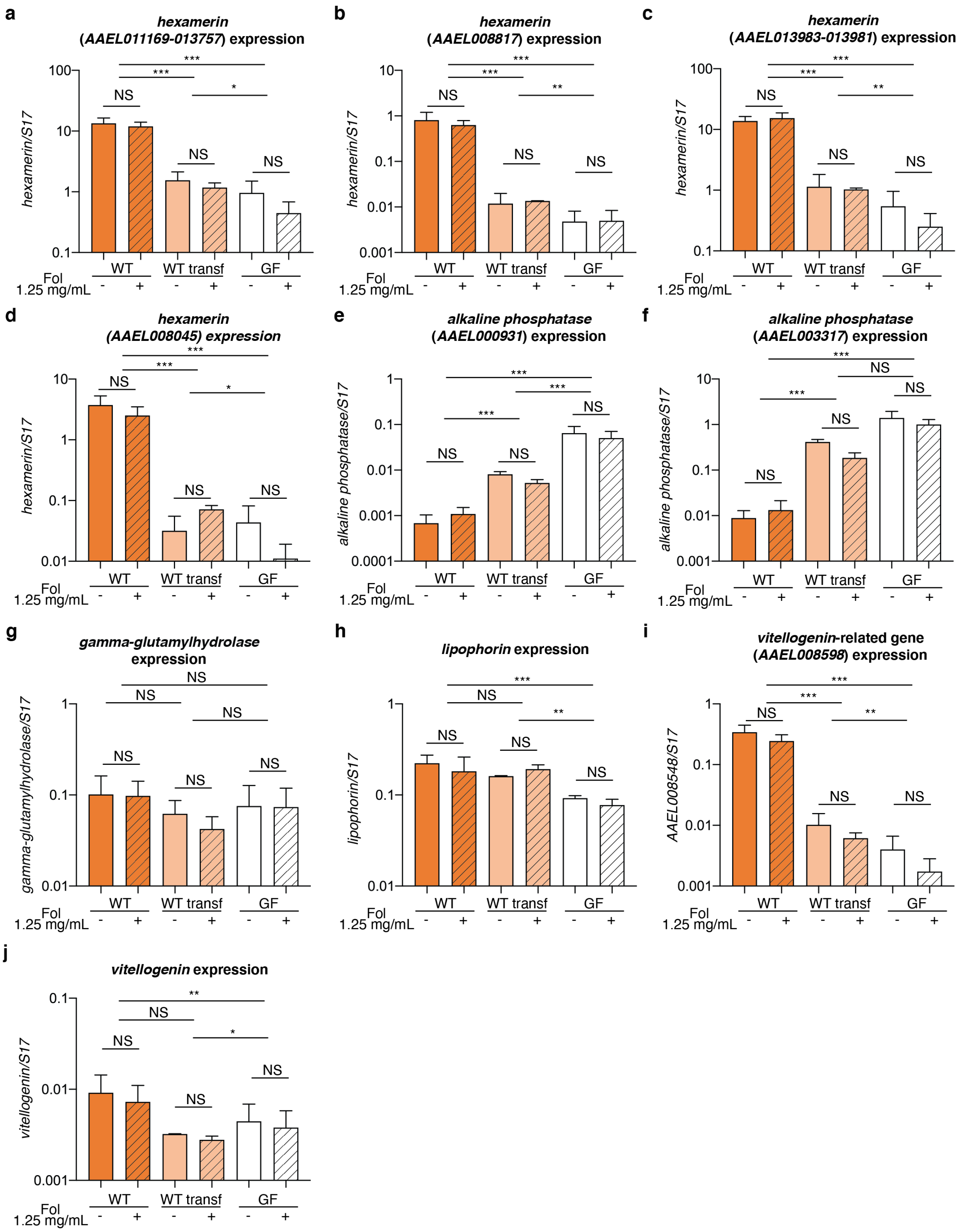


**Figure S13.** Effect of folic acid supplementation (1.25 mg/mL) for 20 h on the expression of *hexamerins AAEL011169-AAEL013757* (**a**), *AAEL008817* (**b**), *AAEL013983-AAEL013981* (**c**), *AAEL008045* (**d**), *alkaline phosphatase AAEL000931* (**e**), *AAEL003317* (**f**), *gamma-glutamyl hydrolase* (**g**), *lipophorin* (**h**), *vitellogenin*-related gene *AAEL008598* (**i**), and *vitellogenin* (**j**) in larvae reared continuously with WT *E. coli* (WT, orange), transferred in new rearing medium after being reared with WT *E. coli* (WT transf, light orange) and after becoming germ-free (GF, white). Data show the mean ± SEM of three independent replicates where at least 18 individuals were analysed per condition, except for expression data on WT transferred larvae which derive from two replicates. Glmm with lsmeans (Bonferroni correction): NS, Non-significant; *, $p < 0.05$, **, $p < 0.01$, ***, $p < 0.001$.

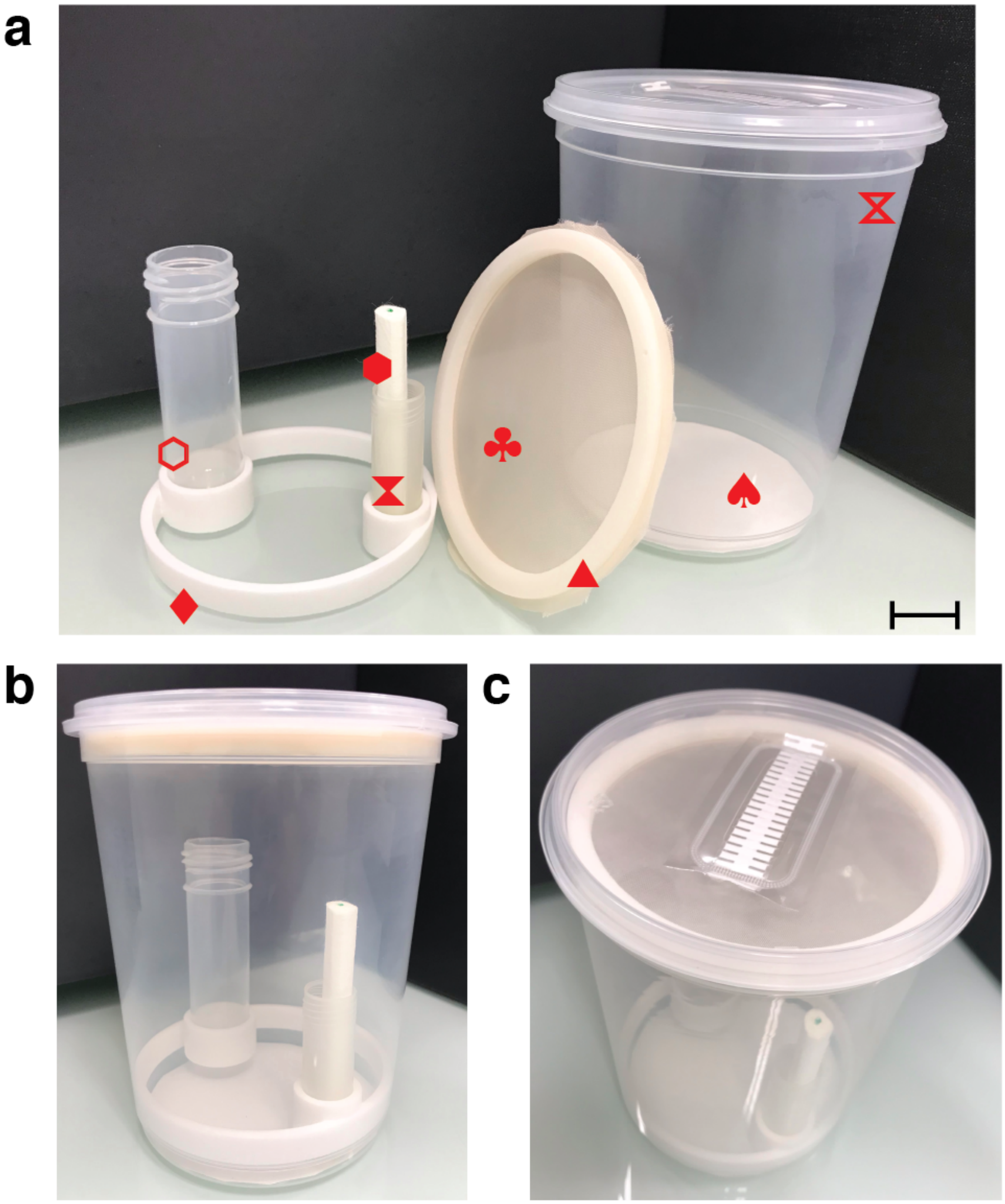

**Figure S14.** Preparation of sterile boxes for germ-free adult mosquitoes. (**a**) Material needed to set-up boxes: autoclavable polypropylene box for plant culture (⧖), larger autoclavable tube (⬡), smaller autoclavable tube (⧗), cotton roll (⬢), mosquito net (♣), filter paper cut to fit the bottom of the box (♠). The filter paper is fixed to the bottom of the box with adhesive tape to collect mosquito excreta. The larger tube will allocate pupae, while in the smaller one a cotton roll is placed for sugar feeding. Boxes are equipped with two components printed in autoclavable material (PA 2200, Shapeways): the first one is designed to hold the two tubes (♦), while the other component consists of two clippable rings holding a disposable mosquito net in the middle (▲). This second component was designed to avoid mosquito contamination through the upper filter and to allow mosquito blood-feeding in sterile conditions. (**b**-**c**) Two different views of the complete box set-up for axenic mosquito rearing. After autoclaving, a sterile 10% sucrose solution is added to the smaller tube with the cotton roll. Scale bar in (**a**) corresponds to 2 cm.

**Table S2.** Number of up- and down-regulated genes in germ-free larvae calculated by DESeq2.

| Number of up- and down-regulated genes in germ-free larvae | | | | | | | |
|---|---|---|---|---|---|---|---|
| | | **Gut 12h** | **Gut 20h** | **Whole larvae 12h** | **Whole larvae 20h** | **Common** | **Total unique genes** |
| Up-regulated genes in germ-free larvae | *padj* < 0.001 | 372 | 614 | 441 | 976 | 121 | 1919 |
| | *padj* < 0.01 | 581 | 910 | 671 | 1312 | 184 | 2738 |
| | *padj* < 0.05 | 864 | 1304 | 929 | 1737 | 269 | 3758 |
| Down-regulated genes in germ-free larvae | *padj* < 0.001 | 295 | 579 | 270 | 580 | 67 | 1456 |
| | *padj* < 0.01 | 493 | 859 | 520 | 966 | 141 | 2274 |
| | *padj* < 0.05 | 766 | 1276 | 881 | 1475 | 240 | 3438 |
| Number of up- and down-regulated genes in germ-free larvae with log2fold values < -1.5 or > 1.5. | | | | | | | |
| | | **Gut 12h** | **Gut 20h** | **Whole larvae 12h** | **Whole larvae 20h** | **Common** | **Total unique genes** |
| Up-regulated genes in germ-free larvae | *padj* < 0.001 | 100 | 203 | 114 | 380 | 25 | 697 |
| | *padj* < 0.01 | 113 | 246 | 134 | 414 | 26 | 803 |
| | *padj* < 0.05 | 130 | 289 | 154 | 472 | 28 | 933 |
| Down-regulated genes in germ-free larvae | *padj* < 0.001 | 99 | 107 | 44 | 138 | 8 | 356 |
| | *padj* < 0.01 | 131 | 125 | 62 | 167 | 10 | 445 |
| | *padj* < 0.05 | 161 | 151 | 85 | 191 | 12 | 540 |

Upper section: Number of up- and down-regulated genes in germ-free larvae. Lower section: Number of up- and down-regulated genes in germ-free larvae with $\log_2$ fold values < -1.5 or > 1.5. In each table, results are shown with several statistical thresholds. "*p*adj" is the adjusted *p* value calculated by DESeq2. "Common" refers to genes up- or down-regulated in all sample types in the same gnotobiotic condition. "Total unique genes" refers to the total number of genes up- or -down-regulated in at least one sample type.

**Table S6.** Expression values of genes encoding for folate transporters.

| AAEL | Description | germ-free vs *E. coli* | germ-free vs *E. coli* | germ-free vs *E. coli* | germ-free vs *E. coli* |
|---|---|---|---|---|---|
| | | GUT_12h | GUT_20h | WL_12h | WL_20h |
| 001687 | Proton-coupled folate transporter | 2.008 | 2.581 | 1.786 | 2.340 |
| 001047 | Proton-coupled folate transporter | 1.775 | 1.268 | 0.874 | 0.494 |
| 001697 | Proton-coupled folate transporter | - | 2.236 | - | - |
| 021114 | Proton-coupled folate transporter | - | - | - | - |
| 001691 | Proton-coupled folate transporter | - | - | - | - |
| 027091 | Folate transporter 1 | - | - | - | - |
| 006879 | Folate carrier protein | - | -0.360 | - | - |
| 003983 | Proton-coupled folate transporter | - | - | 0.782 | -1.064 |
| 009513 | Proton-coupled folate transporter | - | - | - | -1.988 |

For each gene, Vectorbase accession (AAEL), Vectorbase description, and expression values in the different samples are indicated. Significant expression values (*p*adj < 0.001) are coloured with a blue/red colour code corresponding to down/up-regulation. "-" means absence from the DESeq2 output of genes with *p*adj < 0.1, meaning that transcripts are either not detected or not differentially regulated at this 0.1 threshold.

**Table S7.** Primer sequences used in this study.

| Gene name or description | Vectorbase accession | Forward primer | Reverse primer | Reference |
|---|---|---|---|---|
| *16S* | - | TCCTACGGGAGGCAGCAGT | GGACTACCAGGGTATCTAATCCTGTT | [2] |
| *S17* | AAEL004175 | AAGAAGTGGCCATCATTCCA | GGTCTCCGGGTCGACTTC | [3] |
| *Vitellogenin* | AAEL010434 | CTTCTCGCTTTGGCGGGG | CCTGGTAGGCGTTCTGATATCC | - |
| *Hexamerin 2 beta* | AAEL008045 | TCCAAGATGCTGCTCAGTGG | ACGACACCCTTGAAGCTGAG | - |
| *Hexamerin 2 beta* | AAEL008817 | GGTGATCCCAAGTGTCCTGG | CCGGTCGAAGTACGTCACAA | - |
| *Hexamerin 2 beta* | AAEL011169<br>AAEL013757 | CGTACTACTACTACTTCCACGCTG | TCGTTGGACAAGCGTTCCA | - |
| *Hexamerin 2 beta* | AAEL013981<br>AAEL013983 | CTCGTCAGAAGCGAATCAACC | CGACGAAGTCATACACGTATTGATC | - |
| *Unknown (Vitellogenin-related gene)* | AAEL008598 | ACAATTGGGCCGTCTACGTT | CCCGAGACAACTCCATGCTT | - |
| *Lipophorin* | AAEL009955 | GTGGATACCGCGAGTCTCTG | CGTCAGCAGTGGAGTGGATT | - |
| *Alkaline phosphatase* | AAEL000931 | TCGGTTACGCTAATCGACCG | GTAGAAGGCCGATAGGTGCC | - |
| *Alkaline phosphatase* | AAEL003317 | ATGCAGTTACCGGGGATGTC | TTGGTTGGAACCTCGGATGG | - |
| *Gamma-glutamyl hydrolase* | AAEL000271 | ATTGAGACGGCCAAAGGTCC | TCGTTGGCGCATTGAACTTG | - |

For each gene, the Vectorbase accession (if applicable), 5'-3' sequence of the forward and reverse primers and references (if applicable) are indicated.
