## Supplementary material for "Production of germ-free mosquitoes via transient colonisation allows stage-specific investigation of host-microbiota interactions": Table S1

**Table S1.** Statistical information

| Analysis | Response variable | Predictor | Random effect | Sample size | Test result | Comparisons |
|---|---|---|---|---|---|---|
| **Figure 1b. Developmental success** | | | | | | |
| Glmm (binomial) | % adults | Rearing condition | Replicate | Replicate A: AUX (n=76); CR (n=74); WT(n=57)<br>Replicate B: AUX (n=45); CR (n=51); WT (n=48)<br>Replicate C: AUX (n=71); CR (n=62); WT (n=69) | $F_{2,16} = 7.9$, $p < 0.001$ | lsmeans (Bonferroni correction):<br>AUX *vs* CR: $p < 0.001$<br>AUX *vs* WT: $p = 0.87$<br>CR *vs* WT: $p = 0.02$ |
| **Figure S3. Sex ratio** | | | | | | |
| Glmm (binomial) | % females | Rearing condition | Replicate | Replicate A: AUX (n=76); CR (n=74); WT(n=57)<br>Replicate B: AUX (n=45); CR (n=51); WT (n=48)<br>Replicate C: AUX (n=71); CR (n=62); WT (n=69) | $F_{2,8.3} = 4.2$, $p = 0.09$ | / |
| **Figure 1c. Colonisation dynamics in larvae/pupae** | | | | | | |
| Glmm | CFU (L3) day 3 | Bacterium | Replicate | Replicate A: AUX (n=12); WT(n=12)<br>Replicate B: AUX (n=12); WT(n=12)<br>Replicate C: AUX (n=18); WT(n=12) | $F_{1,9e6} = 0.14$, $p = 0.71$ | / |
| Glmm | CFU(L4) day 4 | Bacterium | Replicate | Replicate A: AUX (n=12); WT(n=12)<br>Replicate B: AUX (n=12); WT(n=11)<br>Replicate C: AUX (n=18); WT(n=12) | $F_{1,1e8} = 4.5$, $p = 0.033$ | / |
| Glmm | CFU(L4) day 5 | Bacterium | Replicate | Replicate A: AUX (n=12); WT(n=12)<br>Replicate B: AUX (n=12); WT(n=12)<br>Replicate C: AUX (n=18); WT(n=12) | $F_{1,1e8} = 44$, $p < 0.001$ | / |
| Glmm | CFU(pupae) day 6 | Bacterium | Replicate | Replicate A: AUX (n=12); WT(n=12)<br>Replicate B: AUX (n=12); WT(n=8)<br>Replicate C: AUX (n=13); WT(n=12) | $F_{1,65} = 66$, $p < 0.001$ | / |
| **Figure 1d. Reversible colonisation** | | | | | | |
| Glmm | CFU (adults) day 1 | Bacterium | Replicate | Replicate A: AUX (n=12); WT(n=12)<br>Replicate B: AUX (n=12); WT(n=12)<br>Replicate C: AUX (n=18); WT(n=6) | $F_{1,64} = 5.9$, $p = 0.02$ | / |
| Glmm | CFU (adults) day 2 | Bacterium | Replicate | Replicate A: AUX (n=12); WT(n=12)<br>Replicate B: AUX (n=12); WT(n=12)<br>Replicate C: AUX (n=18); WT(n=6) | $F_{1,64} = 5.1$, $p = 0.03$ | / |
| Glmm | CFU (adults) day 3 | Bacterium | Replicate | Replicate A: AUX (n=12); WT(n=12)<br>Replicate B: AUX (n=12); WT(n=12)<br>Replicate C: AUX (n=18); WT(n=6) | $F_{1,64} = 4.0$, $p = 0.04$ | / |
| **Figure 1e. Bacterial DNA detection via qPCR** | | | | | | |
| | Pools of sugar-fed midguts | | | Replicate A: AUX (n=10); CR (n=10); WT(n=10)<br>Replicate B: AUX (n=10); CR (n=10); WT(n=10)<br>Replicate C: AUX (n=10); CR (n=10); WT(n=10) | | |
| | Pools of blood-fed midguts | | | Replicate A: AUX (n=4); CR (n=5); WT(n=10)<br>Replicate B: AUX (n=6); CR (n=5); WT(n=4)<br>Replicate C: AUX (n=5); CR (n=6); WT(n=4) | | |
| | Pools of sugar-fed whole mosquitoes | | | Replicate A: AUX (n=15); CR (n=12); WT(n=14)<br>Replicate B: AUX (n=12); CR (n=16); WT(n=15)<br>Replicate C: AUX (n=16); CR (n=10); WT(n=16) | | |
| **Figure S4a. Bacterial concentration without larvae** | | | | | | |
| Glmm | CFU (WT) | Time | Replicate | Replicate A: WT (n=1)<br>Replicate B: WT (n=1) | $F_{6,Inf} = 3.4$, $p = 0.002$ | lsmeans (Bonferroni correction):<br>day 0 *vs* day 1: $p = 0.24$ |

| Analysis | Response variable | Predictor | Random effect | Sample size | Test result | Comparisons |
|---|---|---|---|---|---|---|
| | | | | Replicate C: WT (n=1)<br>Replicate D: WT (n=2) | | day 0 *vs* day 2: $p = 0.016$<br>day 0 *vs* day 3: $p = 0.0039$<br>day 0 *vs* day 4: $p = 0.019$<br>day 0 *vs* day 5: $p = 0.044$<br>day 0 *vs* day 6: $p = 1$<br>day 1 *vs* day 2: $p = 1$<br>day 2 *vs* day 3: $p = 1$<br>day 3 *vs* day 4: $p = 1$<br>day 4 *vs* day 5: $p = 1$<br>day 5 *vs* day 6: $p = 1$ |
| Glmm | CFU (AUX) | Time | Replicate | Replicate A: AUX (n=1)<br>Replicate B: AUX (n=1)<br>Replicate C: AUX (n=1)<br>Replicate D: AUX (n=1) | $F_{6,Inf} = 3.0$, $p = 0.006$ | lsmeans (Bonferroni correction):<br>day 0 *vs* day 1: $p = 0.74$<br>day 0 *vs* day 2: $p = 0.028$<br>day 0 *vs* day 3: $p = 0.015$<br>day 0 *vs* day 4: $p = 0.0098$<br>day 0 *vs* day 5: $p = 0.051$<br>day 0 *vs* day 6: $p = 0.30$<br>day 1 *vs* day 2: $p = 1$<br>day 2 *vs* day 3: $p = 1$<br>day 3 *vs* day 4: $p = 1$<br>day 4 *vs* day 5: $p = 1$<br>day 5 *vs* day 6: $p = 1$ |
| Glmm | CFU (day 0) | Bacterium | Replicate | | $F_{1,1e8} = 0.044$, $p = 0.83$ | / |
| Glmm | CFU (day 1) | Bacterium | Replicate | | $F_{1,6e6} = 0.081$, $p = 0.78$ | / |
| Glmm | CFU (day 2) | Bacterium | Replicate | | $F_{1,5802} = 0.84$, $p = 0.36$ | / |
| Glmm | CFU (day 3) | Bacterium | Replicate | | $F_{1,9e4} = 1.8$, $p = 0.18$ | / |
| Glmm | CFU (day 4) | Bacterium | Replicate | | $F_{1,6e47} = 1.2$, $p = 0.27$ | / |
| Glmm | CFU (day 5) | Bacterium | Replicate | | $F_{1,1e5} = 5.7$, $p = 0.017$ | / |
| Glmm | CFU (day 6) | Bacterium | Replicate | | $F_{1,2e4} = 0.85$, $p = 0.35$ | / |
| **Figure S4b. Bacterial concentration with larvae** | | | | | | |
| Glmm | CFU (WT) | Time | Replicate | Replicate A: WT (n=1)<br>Replicate B: WT (n=1)<br>Replicate C: WT (n=1)<br>Replicate D: WT (n=4) | $F_{6,Inf} = 14.16$, $p < 0.001$ | lsmeans (Bonferroni correction):<br>day 0 *vs* day 1: $p = 0.0042$<br>day 0 *vs* day 2: $p < 0.001$<br>day 0 *vs* day 3: $p < 0.001$<br>day 0 *vs* day 4: $p < 0.001$<br>day 0 *vs* day 5: $p < 0.001$<br>day 0 *vs* day 6: $p < 0.001$<br>day 1 *vs* day 2: $p = 1$<br>day 2 *vs* day 3: $p = 1$<br>day 3 *vs* day 4: $p = 0.015$<br>day 4 *vs* day 5: $p = 1$<br>day 5 *vs* day 6: $p = 1$ |
| Glmm | CFU (AUX) | Time | Replicate | Replicate A: AUX (n=1)<br>Replicate B: AUX (n=1)<br>Replicate C: AUX (n=1)<br>Replicate D: AUX (n=5) | $F_{6,Inf} = 16.4$, $p < 0.001$ | lsmeans (Bonferroni correction):<br>day 0 *vs* day 1: $p = 0.0064$<br>day 0 *vs* day 2: $p = 0.098$<br>day 0 *vs* day 3: $p = 0.0090$<br>day 0 *vs* day 4: $p < 0.001$ |

| Analysis | Response variable | Predictor | Random effect | Sample size | Test result | Comparisons |
|---|---|---|---|---|---|---|
| | | | | | | day 0 *vs* day 5: $p < 0.001$<br>day 0 *vs* day 6: $p < 0.001$<br>day 1 *vs* day 2: $p = 1$<br>day 2 *vs* day 3: $p = 1$<br>day 3 *vs* day 4: $p = 0.0074$<br>day 4 *vs* day 5: $p = 1$<br>day 5 *vs* day 6: $p = 1$ |
| Glmm | CFU (day 0) | Bacterium | Replicate | | $F_{1,7e10} = 4.6, p = 0.83$ | / |
| Glmm | CFU (day 1) | Bacterium | Replicate | | $F_{1,2e6} = 0.062, p = 0.80$ | / |
| Glmm | CFU (day 2) | Bacterium | Replicate | | $F_{1,8e5} = 2.4, p = 0.12$ | / |
| Glmm | CFU (day 3) | Bacterium | Replicate | | $F_{1,1e7} = 0.67, p = 0.41$ | / |
| Glmm | CFU (day 4) | Bacterium | Replicate | | $F_{1,2e6} = 0.09, p = 0.76$ | / |
| Glmm | CFU (day 5) | Bacterium | Replicate | | $F_{1,4e9} = 15, p < 0.001$ | / |
| Glmm | CFU (day 6) | Bacterium | Replicate | | $F_{1,5e6} = 39, p < 0.001$ | / |
| **Figure 2a. Duration of development** | | | | | | |
| Glmm | Duration (L1 to L2) | Rearing condition | Replicate | Replicate A: AUX (n=76); CR (n=74); WT(n=57)<br>Replicate B: AUX (n=45); CR (n=51); WT (n=48)<br>Replicate C: AUX (n=71); CR (n=62); WT (n=69) | $F_{2,550} = 18, p < 0.001$ | lsmeans (Bonferroni correction):<br>AUX *vs* CR: $p < 0.001$<br>AUX *vs* WT: $p = 0.0010$<br>CR *vs* WT: $p = 0.095$ |
| Glmm | Duration (L1 to L3) | Rearing condition | Replicate | | $F_{2,526} = 3.1, p = 0.044$ | lsmeans (Bonferroni correction):<br>AUX *vs* CR: $p = 0.13$<br>AUX *vs* WT: $p = 0.07$<br>CR *vs* WT: $p = 1$ |
| Glmm | Duration (L1 to L4) | Rearing condition | Replicate | | $F_{2,503} = 6.9, p = 0.0010$ | lsmeans (Bonferroni correction):<br>AUX *vs* CR: $p = 0.72$<br>AUX *vs* WT: $p < 0.001$<br>CR *vs* WT: $p = 0.051$ |
| Glmm | Duration (L1 to pupa) | Rearing condition | Replicate | | $F_{2,475} = 2.9, p = 0.057$ | / |
| Glmm | Duration (L1 to adult) | Rearing condition | Replicate | | $F_{2,467} = 1.0, p = 0.37$ | / |
| **Figure 2b. Larval length** | | | | | | |
| Glmm | Length | Rearing condition | Replicate | Replicate A: AUX (n=39); CR (n=28); WT(n=55)<br>Replicate B: AUX (n=56); CR (n=32); WT (n=42)<br>Replicate C: AUX (n=30); CR (n=36); WT (n=36)<br>Replicate D: AUX (n=61); CR (n=27); WT (n=45) | $F_{2,481} = 2.5, p = 0.083$ | / |
| **Figure 2c. Wing length** | | | | | | |
| Glmm | Length (females) | Rearing condition | Replicate | Replicate A: AUX (n=11); CR (n=16); WT(n=20)<br>Replicate B: AUX (n=12); CR (n=12); WT (n=7)<br>Replicate C: AUX (n=20); CR (n=8); WT (n=20)<br>Replicate D: AUX (n=20); CR (n=16); WT (n=19) | $F_{2,176} = 1.7, p = 0.19$ | / |
| Glmm | Length (males) | Rearing condition | Replicate | Replicate A: AUX (n=17); CR (n=16); WT(n=20)<br>Replicate B: AUX (n=20); CR (n=15); WT (n=20)<br>Replicate C: AUX (n=8); CR (n=18); WT (n=18)<br>Replicate D: AUX (n=13); CR (n=20); WT (n=20) | $F_{2,199} = 1.9, p = 0.14$ | / |
| **Figure 2d. Egg laying** | | | | | | |

| Analysis | Response variable | Predictor | Random effect | Sample size | Test result | Comparisons |
|---|---|---|---|---|---|---|
| Glmm (binomial) | Egg laying mosquitoes | Rearing condition | Replicate | Replicate A: GF (n=17); CR (n=24); WT(n=25)<br>Replicate B: GF (n=24); CR (n=24); WT (n=14)<br>Replicate C: GF (n=24); CR (n=17); WT (n=12) | $F_{2,27} = 14$, $p < 0.001$ | lsmeans (Bonferroni correction):<br>GF *vs* CR: $p < 0.001$<br>GF *vs* WT: $p = 1$<br>CR *vs* WT: $p < 0.001$ |
| Glmm (binomial) | Egg laying mosquitoes (WT) | Contamination | Replicate | Replicate A: WT cont (n=7); WT non-cont (n=18)<br>Replicate B: WT cont (n=5); WT non-cont (n=9)<br>Replicate C: WT cont (n=6); WT non-cont (n=6) | $F_{2,2} = 1.7$, $p = 0.22$ | / |
| **Figure 2e. Clutch size** | | | | | | |
| Glmm | Number of eggs | Rearing condition | Replicate | See egg laying (Figure 2d) | $F_{2,167} = 9.4$, $p < 0.001$ | lsmeans (Bonferroni correction):<br>GF *vs* CR: $p = 0.0089$<br>GF *vs* WT: $p = 0.78$<br>CR *vs* WT: $p < 0.001$ |
| Glmm | Number of eggs (WT) | Contamination | Replicate | See egg laying (Figure 2d) | $F_{2,48} = 0.11$, $p = 0.74$ | / |
| **Figure S5. Hatching success** | | | | | | |
| Glmm (binomial) | Positive hatching | Rearing condition | Replicate | See egg laying (Figure 2d) | $F_{2,0.1} = 0.062$, $p = 1$ | / |
| Glmm (binomial) | Positive hatching (WT) | Contamination | Replicate | See egg laying (Figure 2d) | $F_{1,0.6} = 0.62$, $p = 0.43$ | / |
| **Figure 2f. Lifespan** | | | | | | |
| Cox model | Survival (females) | Bacterium | / | Replicate A: GF (n=22); WT(n=25)<br>Replicate B: GF (n=21); WT (n=25)<br>Replicate C: GF (n=24); WT (n=19) | z = 1.2, $p = 0.24$<br>0.95 CI (0.87-1.7) | / |
| Cox model | Survival (males) | Bacterium | / | Replicate A: GF (n=28); WT(n=28)<br>Replicate B: GF (n=19); WT (n=22)<br>Replicate C: GF (n=22); WT (n=22) | z = -0.31, $p = 0.76$<br>0.95 CI (0.68-1.3) | / |
| **Figure 3b. Bacterial load after transfer (third instar larvae)** | | | | | | |
| Glmm | CFU (NT) | Bacterium | Replicate | Replicate A: AUX (n=6); WT(n=12)<br>Replicate B: AUX (n=6); WT (n=6)<br>Replicate C: AUX (n=6); WT (n=6) | $F_{1,3e5} = 6.2$, $p = 0.013$ | / |
| Glmm | CFU (2 h) | Bacterium | Replicate | Replicate A: AUX (n=6); WT(n=12)<br>Replicate B: AUX (n=6); WT (n=6)<br>Replicate C: AUX (n=6); WT (n=6) | $F_{1,39} = 0.60$, $p = 0.44$ | / |
| Glmm | CFU (5 h) | Bacterium | Replicate | Replicate A: AUX (n=6); WT(n=11)<br>Replicate B: AUX (n=6); WT (n=6)<br>Replicate C: AUX (n=6); WT (n=6) | $F_{1,37} = 42$, $p < 0.001$ | / |
| Glmm | CFU (12 h) | Bacterium | Replicate | Replicate A: AUX (n=6); WT(n=6)<br>Replicate B: AUX (n=6); WT (n=8)<br>Replicate C: AUX (n=6); WT (n=6) | $F_{1,34} = 19$, $p < 0.001$ | / |
| Glmm | CFU (20 h) | Bacterium | Replicate | Replicate A: AUX (n=6); WT(n=6)<br>Replicate B: AUX (n=6); WT (n=8)<br>Replicate C: AUX (n=6); WT (n=6) | $F_{1,8e4} = 1.2$, $p = 0.26$ | / |
| Glmm | CFU (WT) | Time-point | Replicate | | $F_{4,3e4} = 1.8$, $p = 0.12$ | / |
| Glmm | CFU (AUX) | Time-point | Replicate | | $F_{4,83} = 29$, $p < 0.001$ | lsmeans (Bonferroni correction):<br>NT *vs* 2 h: $p < 0.001$<br>NT *vs* 5 h: $p < 0.001$<br>NT *vs* 12 h: $p < 0.001$ |

| Analysis | Response variable | Predictor | Random effect | Sample size | Test result | Comparisons |
|---|---|---|---|---|---|---|
| | | | | | | NT *vs* 20 h: $p < 0.001$ |
| **Figure 3c. Development success of third-instar larvae** | | | | | | |
| Glmm (binomial) | % L4 | Rearing condition | Replicate | Replicate A: AUX (n=47); WT(n=38); WT transf (n=48)<br>Replicate B: AUX (n=35); WT(n=50); WT transf (n=50)<br>Replicate C: AUX (n=48); WT(n=46); WT transf (n=48)<br>Replicate D: AUX (n=58); WT(n=14); WT transf (n=36)<br>Replicate E: AUX (n=48); WT(n=0); WT transf (n=48)<br>Replicate F: AUX (n=42); WT(n=0); WT transf (n=45)<br>Replicate G: AUX (n=19); WT(n=12); WT transf (n=24) | $F_{2,0.001} = 8*10^{-4}$, $p = 1$ | / |
| Glmm (binomial) | % pupae | Rearing condition | Replicate | | $F_{2,84} = 42$, $p < 0.001$ | lsmeans (Bonferroni correction):<br>GF *vs* WT: $p < 0.001$<br>GF *vs* WT transf: $p = 0.004$<br>WT *vs* WT transf: $p < 0.001$ |
| Glmm (binomial) | % adults | Rearing condition | Replicate | | $F_{2,111} = 55$, $p < 0.001$ | lsmeans (Bonferroni correction):<br>GF *vs* WT: $p < 0.001$<br>GF *vs* WT transf: $p < 0.001$<br>WT *vs* WT transf: $p < 0.001$ |
| **Figure S6. Duration of development** | | | | | | |
| Glmm | Time to pupa | Rearing condition | Replicate | See development success of third-instar larvae (Figure 3c) | $F_{2,370} = 175$, $p < 0.001$ | lsmeans (Bonferroni correction):<br>GF *vs* WT: $p < 0.001$<br>GF *vs* WT transf: $p < 0.001$<br>WT *vs* WT transf: $p < 0.001$ |
| Glmm | Time (L3) | Rearing condition | Replicate | | $F_{2,696} = 16$, $p < 0.001$ | lsmeans (Bonferroni correction):<br>GF *vs* WT: $p < 0.001$<br>GF *vs* WT transf: $p < 0.001$<br>WT *vs* WT transf: $p = 0.75$ |
| Glmm | Time (L4) | Rearing condition | Replicate | | $F_{2,373} = 181$, $p < 0.001$ | lsmeans (Bonferroni correction):<br>GF *vs* WT: $p < 0.001$<br>GF *vs* WT transf: $p < 0.001$<br>WT *vs* WT transf: $p < 0.001$ |
| Glmm | Time (pupal stage) | Rearing condition | Replicate | | $F_{2,345} = 3.3$, $p = 0.039$ | lsmeans (Bonferroni correction):<br>GF *vs* WT: $p = 1$<br>GF *vs* WT transf: $p = 0.055$<br>WT *vs* WT transf: $p = 0.20$ |
| **Figure 3d. Bacterial load 16 h after transfer** | | | | | | |
| Glmm | CFU (transf 24 h) | Bacterium | Replicate | Replicate A: AUX (n=6); WT (n=6)<br>Replicate B: AUX (n=6); WT (n=6)<br>Replicate C: AUX (n=6); WT (n=6) | $F_{1,32} = 20$, $p < 0.001$ | / |
| Glmm | CFU (transf 48 h) | Bacterium | Replicate | Replicate A: AUX (n=6); WT (n=6)<br>Replicate B: AUX (n=6); WT (n=6)<br>Replicate C: AUX (n=6); WT (n=6) | $F_{1,32} = 22$, $p < 0.001$ | / |
| Glmm | CFU (transf 72 h) | Bacterium | Replicate | Replicate A: AUX (n=6); WT (n=6)<br>Replicate B: AUX (n=6); WT (n=6)<br>Replicate C: AUX (n=6); WT (n=6) | $F_{1,34} = 11$, $p = 0.002$ | / |
| Glmm | CFU (transf 96 h) | Bacterium | Replicate | Replicate A: AUX (n=6); WT (n=6)<br>Replicate B: AUX (n=6); WT (n=6)<br>Replicate C: AUX (n=6); WT (n=6) | $F_{1,4e35} = 4.4$, $p = 0.03$ | / |
| **Figure 3e. Development success after transfer at different time-points** | | | | | | |
| Glmm (binomial) | % adults (transf 24 h) | Bacterium | Replicate | Replicate A: AUX (n=44); WT (n=48)<br>Replicate B: AUX (n=37); WT (n=33)<br>Replicate C: AUX (n=47); WT (n=47) | $F_{1,88} = 88$, $p < 0.001$ | / |

| Analysis | Response variable | Predictor | Random effect | Sample size | Test result | Comparisons |
|---|---|---|---|---|---|---|
| | | | | Replicate D: AUX (n=43); WT (n=48) | | |
| Glmm (binomial) | % adults (transf 48 h) | Bacterium | Replicate | Replicate A: AUX (n=47); WT (n=48)<br>Replicate B: AUX (n=36); WT (n=34)<br>Replicate C: AUX (n=45); WT (n=48)<br>Replicate D: AUX (n=48); WT (n=47) | $F_{1,67} = 67$, $p < 0.001$ | / |
| Glmm (binomial) | % adults (transf 72 h) | Bacterium | Replicate | Replicate A: AUX (n=48); WT (n=47)<br>Replicate B: AUX (n=46); WT (n=46)<br>Replicate C: AUX (n=45); WT (n=44)<br>Replicate D: AUX (n=46); WT (n=46) | $F_{1,11} = 11$, $p < 0.001$ | / |
| Glmm (binomial) | % adults (transf 96 h) | Bacterium | Replicate | Replicate A: AUX (n=48); WT (n=44)<br>Replicate B: AUX (n=46); WT (n=44)<br>Replicate C: AUX (n=48); WT (n=46)<br>Replicate D: AUX (n=41); WT (n=42) | $F_{1,0.5} = 0.49$, $p = 0.48$ | / |
| Glmm (binomial) | % dead *vs* blocked (transf 24 h) | Bacterium | Replicate | See % adults (transf 24 h) | $F_{1,13} = 13$, $p < 0.001$ | / |
| Glmm (binomial) | % dead *vs* blocked (transf 48 h) | Bacterium | Replicate | See % adults (transf 48 h) | $F_{1,1} = 1.2$, $p = 0.28$ | / |
| Glmm (binomial) | % dead *vs* blocked (transf 72 h) | Bacterium | Replicate | See % adults (transf 72 h) | $F_{1,2} = 2.1$, $p = 0.18$ | / |
| Glmm (binomial) | % dead *vs* blocked (transf 96 h) | Bacterium | Replicate | See % adults (transf 96 h) | $F_{1,0.4} = 0.41$, $p = 0.53$ | / |
| **Figure S7. Duration of development after transfer** | | | | | | |
| Glmm | Time to pupation (transf 24 h) | Bacterium | Replicate | See % adults (transf 24 h, Figure 3e) | $F_{1,143} = 11$, $p < 0.001$ | / |
| Glmm | Time to pupation (transf 48 h) | Bacterium | Replicate | See % adults (transf 48 h, Figure 3e) | $F_{1,116} = 67$, $p < 0.001$ | / |
| Glmm | Time to pupation (transf 72 h) | Bacterium | Replicate | See % adults (transf 72 h, Figure 3e) | $F_{1,279} = 24$, $p < 0.001$ | / |
| Glmm | Time to pupation (transf 96 h) | Bacterium | Replicate | See % adults (transf 96 h, Figure 3e) | $F_{1,330} = 13$, $p < 0.001$ | / |
| **Figure 4a. CFU after transfer (breeding site water)** | | | | | | |
| Glmm | CFU NT *vs* 2-5 h (transf 48 h) | Transfer | Replicate | Replicate A: NT (n=6); TR-2h (n=5); TR-5h (n=6)<br>Replicate B: NT (n=6); TR-2h (n=6); TR-5h (n=6) | $F_{2,31} = 9.7$, $p < 0.001$ | lsmeans (Bonferroni correction):<br>NT *vs* 2 h: $p = 0.0020$<br>NT *vs* 5 h: $p = 0.0018$<br>2 h *vs* 5 h: $p = 1$ |
| Glmm | CFU NT-24 h *vs* 24 h (transf 48 h) | Transfer | Replicate | Replicate A: NT-24h (n=6); TR-24h (n=6)<br>Replicate B: NT-24h (n=6); TR-24h (n=6) | $F_{1,2e20} = 0.85$, $p = 0.36$ | / |
| **Figure 4c. CFU after transfer (breeding site water)** | | | | | | |
| Glmm | CFU NT *vs* 2-5 h (transf 72 h) | Transfer | Replicate | Replicate A: NT (n=6); TR-2h (n=6); TR-5h (n=6)<br>Replicate B: NT (n=6); TR-2h (n=6); TR-5h (n=6)<br>Replicate C: NT (n=6); TR-2h (n=6); TR-5h (n=6) | $F_{2,49} = 20$, $p < 0.001$ | lsmeans (Bonferroni correction):<br>NT *vs* 2 h: $p < 0.001$<br>NT *vs* 5 h: $p < 0.001$<br>2 h *vs* 5 h: $p = 1$ |
| Glmm | CFU NT-24 h *vs* 24 h (transf 72 h) | Transfer | Replicate | Replicate A: NT-24h (n=6); TR-24h (n=6)<br>Replicate B: NT-24h (n=6); TR-24h (n=6)<br>Replicate C: NT-24h (n=6); TR-24h (n=6) | $F_{1,3e5} = 1.7$, $p = 0.18$ | / |
| **Figure 4b. Development success after transfer (breeding site water)** | | | | | | |

| Analysis | Response variable | Predictor | Random effect | Sample size | Test result | Comparisons |
|---|---|---|---|---|---|---|
| Glmm (binomial) | % adults (transf 48 h) | Transfer | Replicate | Replicate A: NT (n=44); TR-48 (n=44)<br>Replicate B: NT (n=40); TR-48 (n=39)<br>Replicate C: NT (n=50); TR-48 (n=44) | $F_{1,14} = 14$, $p < 0.001$ | / |
| Glmm (binomial) | % dead *vs* blocked (transf 48 h) | Transfer | Replicate | | $F_{1,17} = 17$, $p = 0.0081$ | / |
| **Figure 4d. Development success after transfer (breeding site water)** | | | | | | |
| Glmm (binomial) | % adults (transf 72 h) | Transfer | Replicate | Replicate A: NT (n=40); TR-72 (n=43)<br>Replicate B: NT (n=43); TR-72 (n=44)<br>Replicate C: NT (n=36); TR-72 (n=37)<br>Replicate D: NT (n=44); TR-72 (n=46)<br>Replicate E: NT (n=29); TR-72 (n=39)<br>Replicate F: NT (n=40); TR-72 (n=45)<br>Replicate G: NT (n=50); TR-72 (n=46) | $F_{1,9} = 9.3$, $p = 0.002$ | / |
| Glmm (binomial) | % dead *vs* blocked (transf 72 h) | Transfer | Replicate | | $F_{1,22} = 22$, $p < 0.001$ | / |
| **Figure S8. *16S* qPCR on sequenced samples** | | | | | | |
| Glmm | $Log_2$ Ratio (gut 12 h) | Bacterium | Replicate | Replicate A: GF (n=1, pool of 60); WT (n=1, pool of 60)<br>Replicate B: GF (n=1, pool of 60); WT (n=1, pool of 60)<br>Replicate C: GF (n=1, pool of 60); WT (n=1, pool of 60) | $F_{1,2} = 92$, $p = 0.011$ | / |
| Glmm | $Log_2$ Ratio (gut 20 h) | Bacterium | Replicate | | $F_{1,4} = 631$, $p < 0.001$ | / |
| Glmm | $Log_2$ Ratio (wl 12 h) | Bacterium | Replicate | | $F_{1,4} = 114$, $p < 0.001$ | / |
| Glmm | $Log_2$ Ratio (wl 20 h) | Bacterium | Replicate | | $F_{1,2} = 2704$, $p < 0.001$ | / |
| **Figure 6b. Development success with folate** | | | | | | |
| Glmm (binomial) | % L4 | Folate concentration | Replicate | Replicate A:<br>GF (n=48); 0.25 FOL (n=48); 0.5 FOL (n=46); 1.25 FOL (n=48)<br>Replicate B:<br>GF (n=42); 0.25 FOL (n=24); 0.5 FOL (n=24); 1.25 FOL (n=24)<br>Replicate C:<br>GF (n=19); 0.25 FOL (n=19); 0.5 FOL (n=19); 1.25 FOL (n=20) | $F_{3,7e\text{-}11} = 0$, $p = 1$ | / |
| Glmm (binomial) | % pupae | Folate concentration | Replicate | | $F_{3,51} = 17$, $p < 0.001$ | lsmeans (Bonferroni correction):<br>GF *vs* 0.25 FOL: $p < 0.001$<br>GF *vs* 0.5 FOL: $p < 0.001$<br>GF *vs* 1.25 FOL: $p < 0.00$<br>All other comparisons: $p > 0.05$ |
| Glmm (binomial) | % adults | Folate concentration | Replicate | | $F_{3,52} = 17$, $p < 0.001$ | lsmeans (Bonferroni correction):<br>GF *vs* 0.25 FOL: $p < 0.001$<br>GF *vs* 0.5 FOL: $p < 0.001$<br>GF *vs* 1.25 FOL: $p < 0.00$<br>All other comparisons: $p > 0.05$ |
| **Figure S11. Duration of development with folate** | | | | | | |
| Glmm | Time to pupa | Folate concentration | Replicate | Replicate A:GF (n=48); 0.25 FOL (n=48); 0.5 FOL (n=46); 1.25 FOL (n=48)<br>Replicate B: GF (n=42); 0.25 FOL (n=24); 0.5 FOL (n=24); 1.25 FOL (n=24)<br>Replicate C: GF (n=19); 0.25 FOL (n=19); 0.5 FOL (n=19); 1.25 FOL (n=20) | $F_{3,151} = 1.4$, $p = 0.26$ | / |
| Glmm | Time (L3) | Folate concentration | Replicate | | $F_{3,376} = 0.68$, $p = 0.57$ | / |
| Glmm | Time (L4) | Folate concentration | Replicate | | $F_{3,162} = 1.4$, $p = 0.26$ | / |
| Glmm | Time (pupal stage) | Folate concentration | Replicate | | $F_{3,152} = 1.1$, $p = 0.34$ | / |
| **Figure S12. Effect of folate on L1 axenic larvae** | | | | | | |
| | | | | WT (n=48); GF+FOL (n=24) | | |
| **Figure 6d. Gut lipid quantification** | | | | | | |
| Glmm | Fluorescence/Area | Rearing condition | Replicate | Replicate A: GF (n=12); WT(n=9); WT transf (n=14)<br>Replicate B: GF (n=19); WT(n=18); WT transf (n=13)<br>Replicate C: GF (n=13); WT(n=16); WT transf (n=15) | $F_{2,124} = 7.7$, $p < 0.001$ | lsmeans (Bonferroni correction):<br>GF *vs* WT: $p = 0.0026$<br>GF *vs* WT transf: $p = 0.0033$<br>WT *vs* WT transf: $p = 1$ |

| Analysis | Response variable | Predictor | Random effect | Sample size | Test result | Comparisons |
|---|---|---|---|---|---|---|
| **Figure 6e. Pelt lipid quantification** | | | | | | |
| Glmm | Fluorescence | Rearing condition | Replicate | Replicate A: GF (n=12); WT(n=12); WT transf (n=12)<br>Replicate B: GF (n=12); WT(n=12); WT transf (n=12)<br>Replicate C: GF (n=10); WT(n=10); WT transf (n=10) | $F_{2,97}$ = 18, $p$ < 0.001 | lsmeans (Bonferroni correction):<br>GF *vs* WT: $p$ < 0.001<br>GF *vs* WT transf: $p$ = 0.0045<br>WT *vs* WT transf: $p$ = 0.019 |
| **Figure 6f. Pelt DNA quantification** | | | | | | |
| Glmm | Fluorescence | Rearing condition | Replicate | Replicate A: GF (n=12); WT(n=12); WT transf (n=12)<br>Replicate B: GF (n=12); WT(n=12); WT transf (n=12)<br>Replicate C: GF (n=10); WT(n=10); WT transf (n=10) | $F_{2,97}$ = 16, $p$ < 0.001 | lsmeans (Bonferroni correction):<br>GF *vs* WT: $p$ < 0.001<br>GF *vs* WT transf: $p$ = 0.0019<br>WT *vs* WT transf: $p$ = 0.12 |
| **Figure 6g. Larval length** | | | | | | |
| Glmm | Length | Rearing condition | Replicate | Replicate A: GF (n=10); WT(n=9); WT transf (n=9)<br>Replicate B: GF (n=24); WT(n=24); WT transf (n=24)<br>Replicate C: GF (n=12); WT(n=6); WT transf (n=12) | $F_{2,125}$ = 18, $p$ < 0.001 | lsmeans (Bonferroni correction):<br>GF *vs* WT: $p$ = 0.001<br>GF *vs* WT transf: $p$ < 0.001<br>WT *vs* WT transf: $p$ = 0.77 |
| **Figure S10b. Hypoxia measurements** | | | | | | |
| Glmm | Fluorescence | Rearing condition | Replicate | Replicate A: GF (n=10); WT(n=10); WT transf (n=10)<br>Replicate B: GF (n=10); WT(n=10); WT transf (n=10)<br>Replicate C: GF (n=9); WT(n=9); WT transf (n=9) | $F_{2,82}$ = 0.52, $p$ = 0.60 | / |
| **Figure S13. qPCR** | | | | | | |
| Glmm | $Log_2$ Ratio *Hexamerin 2 beta* (*AAEL011169*, *AAEL013757*) | Folate addition | Replicate | Replicate A:<br>GF (n=1, pool of 24); GF+FOL (n=1, pool of 24);<br>WT (n=1, pool of 18); WT+FOL (n=1, pool of 18);<br>WT transf (n=1, pool of 24); WT transf + FOL (n=1, pool of 24)<br><br>Replicate B:<br>GF (n=1, pool of 24); GF+FOL (n=1, pool of 24);<br>WT (n=1, pool of 22); WT+FOL (n=1, pool of 22);<br>WT transf (n=1, pool of 24); WT transf + FOL (n=1, pool of 24)<br><br>Replicate C:<br>GF (n=1, pool of 24); GF+FOL (n=1, pool of 24);<br>WT (n=1, pool of 24); WT+FOL (n=1, pool of 24);<br>WT transf (n=0); WT transf + FOL (n=0) | $F_{5,8}$ = 23, $p$ < 0.001 | lsmeans (Bonferroni correction):<br>WT *vs* WT-FOL: $p$ = 1<br>WT transf *vs* WT transf-FOL: $p$ = 1<br>GF *vs* GF-FOL: $p$ = 1 |
| Glmm | $Log_2$ Ratio *Hexamerin 2 beta* (*AAEL011169*, *AAEL013757*) | Rearing condition | Replicate | | $F_{2,11}$ = 55, $p$ < 0.001 | lsmeans (Bonferroni correction):<br>GF *vs* WT: $p$ < 0.001<br>GF *vs* WT transf: $p$ = 0.022<br>WT *vs* WT transf: $p$ < 0.001 |
| Glmm | $Log_2$ Ratio *Hexamerin 2 beta* (*AAEL008817*) | Folate addition | Replicate | | $F_{5,8}$ = 45, $p$ < 0.001 | lsmeans (Bonferroni correction):<br>WT *vs* WT-FOL: $p$ = 1<br>WT transf *vs* WT transf-FOL: $p$ = 1<br>GF *vs* GF-FOL: $p$ = 1 |
| Glmm | $Log_2$ Ratio *Hexamerin 2 beta* (*AAEL008817*) | Rearing condition | Replicate | | $F_{2,11}$ = 144, $p$ < 0.001 | lsmeans (Bonferroni correction):<br>GF *vs* WT: $p$ < 0.001<br>GF *vs* WT transf: $p$ = 0.002<br>WT *vs* WT transf: $p$ < 0.001 |
| Glmm | $Log_2$ Ratio *Hexamerin 2 beta* (*AAEL013981*, *AAEL013983*) | Folate addition | Replicate | | $F_{5,8}$ = 18, $p$ < 0.001 | lsmeans (Bonferroni correction):<br>WT *vs* WT-FOL: $p$ = 1<br>WT transf *vs* WT transf-FOL: $p$ = 1<br>GF *vs* GF-FOL: $p$ = 1 |
| Glmm | $Log_2$ Ratio *Hexamerin 2 beta* (*AAEL013981*, *AAEL013983*) | Rearing condition | Replicate | | $F_{2,11}$ = 11, $p$ < 0.001 | lsmeans (Bonferroni correction):<br>GF *vs* WT: $p$ < 0.001<br>GF *vs* WT transf: $p$ = 0.005<br>WT *vs* WT transf: $p$ < 0.001 |
| Glmm | $Log_2$ Ratio *Hexamerin 2 beta* (*AAEL008045*) | Folate addition | Replicate | | $F_{5,8}$ = 22, $p$ < 0.001 | lsmeans (Bonferroni correction):<br>WT *vs* WT-FOL: $p$ = 1 |

| Analysis | Response variable | Predictor | Random effect | Sample size | Test result | Comparisons |
|---|---|---|---|---|---|---|
| | | | | | | WT transf *vs* WT transf-FOL: $p = 1$<br>GF *vs* GF-FOL: $p = 1$ |
| Glmm | $Log_2$ Ratio *Hexamerin 2 beta* (*AAEL008045*) | Rearing condition | Replicate | | $F_{2,11} = 51, p < 0.001$ | lsmeans (Bonferroni correction):<br>GF *vs* WT: $p < 0.001$<br>GF *vs* WT transf: $p = 0.038$<br>WT *vs* WT transf: $p < 0.001$ |
| Glmm | $Log_2$ Ratio *Alkaline phosphatase* (*AAEL000931*) | Folate addition | Replicate | | $F_{5,8} = 108, p < 0.001$ | lsmeans (Bonferroni correction):<br>WT *vs* WT-FOL: $p = 0.78$<br>WT transf *vs* WT transf-FOL: $p = 1$<br>GF *vs* GF-FOL: $p = 1$ |
| Glmm | $Log_2$ Ratio *Alkaline phosphatase* (*AAEL000931*) | Rearing condition | Replicate | | $F_{2,11} = 183, p < 0.001$ | lsmeans (Bonferroni correction):<br>GF *vs* WT: $p < 0.001$<br>GF *vs* WT transf: $p < 0.001$<br>WT *vs* WT transf: $p < 0.001$ |
| Glmm | $Log_2$ Ratio *Alkaline phosphatase* (*AAEL003317*) | Folate addition | Replicate | | $F_{5,10} = 17, p < 0.001$ | lsmeans (Bonferroni correction):<br>WT *vs* WT-FOL: $p = 1$<br>WT transf *vs* WT transf-FOL: $p = 1$<br>GF *vs* GF-FOL: $p = 1$ |
| Glmm | $Log_2$ Ratio *Alkaline phosphatase* (*AAEL003317*) | Rearing condition | Replicate | | $F_{2,13} = 51, p < 0.001$ | lsmeans (Bonferroni correction):<br>GF *vs* WT: $p < 0.001$<br>GF *vs* WT transf: $p = 0.10$<br>WT *vs* WT transf: $p < 0.001$ |
| Glmm | $Log_2$ Ratio *Gamma-glutamyl hydrolase* (*AAEL000271*) | Folate addition | Replicate | | $F_{5,8} = 1.9, p = 0.21$ | / |
| Glmm | $Log_2$ Ratio *Gamma-glutamyl hydrolase* (*AAEL000271*) | Rearing condition | Replicate | | $F_{2,11} = 4.9, p = 0.029$ | lsmeans (Bonferroni correction):<br>GF *vs* WT: $p = 0.077$<br>GF *vs* WT transf: $p = 0.059$<br>WT *vs* WT transf: $p = 1$ |
| Glmm | $Log_2$ Ratio *Lipophorin* (*AAEL009955*) | Folate addition | Replicate | | $F_{5,8} = 9.9, p = 0.0028$ | lsmeans (Bonferroni correction):<br>WT *vs* WT-FOL: $p = 1$<br>WT transf *vs* WT transf-FOL: $p = 1$<br>GF *vs* GF-FOL: $p = 1$ |
| Glmm | $Log_2$ Ratio *Lipophorin* (*AAEL009955*) | Rearing condition | Replicate | | $F_{2,11} = 27, p < 0.001$ | lsmeans (Bonferroni correction):<br>GF *vs* WT: $p < 0.001$<br>GF *vs* WT transf: $p = 0.0042$<br>WT *vs* WT transf: $p = 0.21$ |
| Glmm | $Log_2$ Ratio *Vitellogenin*-related gene (*AAEL008598*) | Folate addition | Replicate | | $F_{5,8} = 44, p < 0.001$ | lsmeans (Bonferroni correction):<br>WT *vs* WT-FOL: $p = 1$<br>WT transf *vs* WT transf-FOL: $p = 1$<br>GF *vs* GF-FOL: $p = 1$ |
| Glmm | $Log_2$ Ratio *Vitellogenin*-related gene (*AAEL008598*) | Rearing condition | Replicate | | $F_{2,11} = 106, p < 0.001$ | lsmeans (Bonferroni correction):<br>GF *vs* WT: $p < 0.001$<br>GF *vs* WT transf: $p = 0.0031$<br>WT *vs* WT transf: $p < 0.001$ |
| Glmm | $Log_2$ Ratio *Vitellogenin* (*AAEL010434*) | Folate addition | Replicate | | $F_{5,8} = 5.1, p = 0.02$ | lsmeans (Bonferroni correction):<br>WT *vs* WT-FOL: $p = 1$<br>WT transf *vs* WT transf-FOL: $p = 1$ |

| Analysis | Response variable | Predictor | Random effect | Sample size | Test result | Comparisons |
|---|---|---|---|---|---|---|
| | | | | | | GF *vs* GF-FOL: $p = 1$ |
| Glmm | $Log_2$ Ratio *Vitellogenin* (*AAEL010434*) | Rearing condition | Replicate | | $F_{2,11} = 13$, $p = 0.001$ | lsmeans (Bonferroni correction):<br>GF *vs* WT: $p = 0.001$<br>GF *vs* WT transf: $p = 0.038$<br>WT *vs* WT transf: $p = 0.70$ |

For each figure, the type of analysis, response variable (dependent variable), predictor (independent variable), random effect, sample size in each replicate, test results and multiple comparisons (if applicable) are shown. Glmm: generalised linear mixed model. lsmeans: least square means.
