## Supplementary material for "Production of germ-free mosquitoes via transient colonisation allows stage-specific investigation of host-microbiota interactions": Table S3

**Table S3**. Gene Ontology (GO) and KEGG terms significantly enriched in up-regulated (red) and down-regulated (blue) genes in germ-free larvae compared to colonised larvae.

| Sample | Source | Term name | Term ID | Adjusted *p* value | n |
|---|---|---|---|---|---|
| Gut, 12H | **GO:MF** | oxidoreductase activity, acting on paired donors, with incorporation or reduction of molecular oxygen | GO:0016705 | 8.5E-07 | 12 |
| | **GO:MF** | heme binding | GO:0020037 | 8.9E-07 | 12 |
| | **GO:MF** | tetrapyrrole binding | GO:0046906 | 9.4E-07 | 12 |
| | **GO:MF** | monooxygenase activity | GO:0004497 | 1.6E-06 | 11 |
| | **GO:MF** | cofactor binding | GO:0048037 | 1.1E-05 | 15 |
| | **GO:MF** | iron ion binding | GO:0005506 | 1.3E-05 | 11 |
| | **GO:MF** | alkaline phosphatase activity | GO:0004035 | 2.1E-05 | 4 |
| | **GO:MF** | catalytic activity | GO:0003824 | 4.5E-05 | 42 |
| | **GO:MF** | oxidoreductase activity | GO:0016491 | 2.1E-03 | 15 |
| | **GO:MF** | phosphatase activity | GO:0016791 | 6.4E-03 | 6 |
| | **GO:MF** | phosphoric ester hydrolase activity | GO:0042578 | 3.0E-02 | 6 |
| | **GO:BP** | oxidation-reduction process | GO:0055114 | 1.5E-02 | 16 |
| | **KEGG** | Folate biosynthesis | KEGG:00790 | 1.9E-03 | 3 |
| | **KEGG** | Thiamine metabolism | KEGG:00730 | 7.3E-03 | 2 |
| | **GO:MF** | phospholipase A1 activity | GO:0008970 | 3.6E-02 | 2 |
| Gut, 20H | **GO:MF** | alkaline phosphatase activity | GO:0004035 | 2.1E-04 | 4 |
| | **GO:MF** | cofactor binding | GO:0048037 | 3.8E-03 | 16 |
| | **GO:MF** | oxidoreductase activity, acting on paired donors, with incorporation or reduction of molecular oxygen | GO:0016705 | 1.5E-02 | 10 |
| | **GO:MF** | heme binding | GO:0020037 | 1.6E-02 | 10 |
| | **GO:MF** | tetrapyrrole binding | GO:0046906 | 1.6E-02 | 10 |
| | **GO:MF** | iron ion binding | GO:0005506 | 1.9E-02 | 10 |
| | **GO:MF** | catalytic activity | GO:0003824 | 2.0E-02 | 58 |
| | **GO:MF** | monooxygenase activity | GO:0004497 | 2.0E-02 | 9 |
| | **GO:MF** | metallocarboxypeptidase activity | GO:0004181 | 3.4E-02 | 4 |
| | **GO:CC** | extracellular region | GO:0005576 | 8.4E-04 | 12 |
| | **KEGG** | Steroid biosynthesis | KEGG:00100 | 1.6E-02 | 2 |
| | **KEGG** | Thiamine metabolism | KEGG:00730 | 3.9E-02 | 2 |
| Whole larvae, 12h | **GO:MF** | catalytic activity | GO:0003824 | 1.3E-05 | 46 |
| | **GO:MF** | alkaline phosphatase activity | GO:0004035 | 4.1E-05 | 4 |
| | **GO:MF** | heme binding | GO:0020037 | 3.2E-04 | 10 |
| | **GO:MF** | tetrapyrrole binding | GO:0046906 | 3.4E-04 | 10 |
| | **GO:MF** | oxidoreductase activity, acting on paired donors, with incorporation or reduction of molecular oxygen | GO:0016705 | 2.6E-03 | 9 |
| | **GO:MF** | iron ion binding | GO:0005506 | 3.2E-03 | 9 |
| | **GO:MF** | monooxygenase activity | GO:0004497 | 5.4E-03 | 8 |
| | **GO:MF** | phosphatase activity | GO:0016791 | 1.4E-02 | 6 |
| | **GO:MF** | structural constituent of cuticle | GO:0042302 | 4.0E-02 | 8 |
| | **GO:MF** | cofactor binding | GO:0048037 | 4.3E-02 | 11 |
| | **KEGG** | Thiamine metabolism | KEGG:00730 | 5.0E-04 | 2 |
| | **KEGG** | Folate biosynthesis | KEGG:00790 | 5.7E-03 | 2 |
| | **GO:MF** | structural constituent of cuticle | GO:0042302 | 7.6E-04 | 7 |
| | **GO:MF** | structural molecule activity | GO:0005198 | 3.0E-02 | 7 |
| | **KEGG** | Steroid biosynthesis | KEGG:00100 | 5.0E-02 | 1 |
| Whole larvae, 20h | **GO:MF** | structural constituent of cuticle | GO:0042302 | 4.8E-08 | 21 |
| | **GO:MF** | structural molecule activity | GO:0005198 | 9.5E-04 | 21 |
| | **GO:MF** | chitin binding | GO:0008061 | 1.1E-03 | 12 |
| | **GO:MF** | alkaline phosphatase activity | GO:0004035 | 1.4E-03 | 4 |
| | **GO:MF** | endopeptidase activity | GO:0004175 | 3.0E-02 | 20 |
| | **GO:MF** | peptidase activity, acting on L-amino acid peptides | GO:0070011 | 4.4E-02 | 24 |
| | **GO:BP** | chitin metabolic process | GO:0006030 | 4.6E-05 | 12 |
| | **GO:BP** | glucosamine-containing compound metabolic process | GO:1901071 | 5.5E-05 | 12 |
| | **GO:BP** | amino sugar metabolic process | GO:0006040 | 6.4E-05 | 12 |
| | **GO:BP** | aminoglycan metabolic process | GO:0006022 | 1.4E-04 | 12 |
| | **GO:BP** | metabolic process | GO:0008152 | 2.1E-04 | 79 |
| | **GO:BP** | proteolysis | GO:0006508 | 9.0E-04 | 27 |
| | **GO:BP** | drug metabolic process | GO:0017144 | 5.1E-03 | 12 |
| | **GO:CC** | extracellular region | GO:0005576 | 2.1E-07 | 20 |
| | **KEGG** | Thiamine metabolism | KEGG:00730 | 9.8E-03 | 2 |
| | **GO:MF** | serine-type peptidase activity | GO:0008236 | 2.2E-03 | 12 |
| | **GO:MF** | serine hydrolase activity | GO:0017171 | 2.2E-03 | 12 |
| | **GO:MF** | serine-type endopeptidase activity | GO:0004252 | 6.4E-03 | 11 |
| | **GO:MF** | oxidoreductase activity, acting on the aldehyde or oxo group of donors, NAD or NADP as acceptor | GO:0016620 | 3.9E-02 | 4 |
| | **KEGG** | Tyrosine metabolism | KEGG:00350 | 3.7E-02 | 2 |
| | **KEGG** | Metabolic pathways | KEGG:01100 | 4.0E-02 | 7 |

The analysis was performed in g:Profiler. Statistical significance was assessed with a Fisher's one tail test with Bonferroni correction. "n" indicates the number of genes associated with that term that are significanlty regulated in the indicated sample. GO:MF: Gene Ontology, Molecular Function; GO:BP: Gene Ontology, Biological Process; GO:CC: Gene Ontology, Cell Compartment.
