## Supplementary material for "Production of germ-free mosquitoes via transient colonisation allows stage-specific investigation of host-microbiota interactions": Table S4

**Table S4**. List of the most up-regulated genes in germ-free larvae compared to colonised larvae.

| AAEL | Description | Phosphatase activity | Hydrolytic reaction | Chitin binding | Transferase activity | Immunity | Cuticle constituent | Lipid metabolism | Other functions | germ-free vs *E. coli* GUT_12h | germ-free vs *E. coli* GUT_20h | germ-free vs *E. coli* WL_12h | germ-free vs *E. coli* WL_20h |
|---|---|---|---|---|---|---|---|---|---|---|---|---|---|
| 000931 | Alkaline phosphatase | | | | | | | | | 9.2 | 7.2 | 5.5 | 5.6 |
| 003317 | Alkaline phosphatase | | | | | | | | | 7.1 | 6.1 | 7.6 | 6.0 |
| 003297 | Alkaline phosphatase | | | | | | | | | 5.8 | 3.2 | 5.4 | 3.2 |
| 002555 | Sodium/solute symporter | | | | | | | | Transmembrane transport | 5.7 | 4.5 | 3.6 | 3.1 |
| 002138 | Triacylglycerol lipase | | | | | | | | | 4.4 | 3.9 | 4.0 | 4.0 |
| 003289 | Alkaline phosphatase | | | | | | | | | 5.2 | 3.0 | 4.7 | 3.1 |
| 000323 | Cysteine-rich venom protein | | | | | | | | Trypsine inhibitor activity | 4.0 | 3.9 | 3.7 | 3.6 |
| 029062 | Unknown (Chitin binding) | | | | | | | | | 5.0 | 3.0 | 4.5 | 2.7 |
| 024870 | Nose resistant to fluoxetine protein 6-like | | | | | | | | | 3.3 | 3.1 | 3.8 | 3.1 |
| 003905 | Alkaline phosphatase | | | | | | | | | 4.5 | 2.5 | 3.1 | 3.1 |
| 012636 | Cytochrome b5 | | | | | | | | Oxidoreductase activity | 3.0 | 3.5 | 3.0 | 2.7 |
| 014246 | Glucosyl/glucuronosyl transferases | | | | | | | | | 4.1 | 2.7 | 3.7 | 2.6 |
| 017359 | Unknown | | | | | | | | | -1.3 | 5.8 | - | 4.4 |
| 012955 | Phosphatidylethanolamine-binding protein | | | | | | | | | 3.4 | 3.7 | 2.0 | 1.1 |
| 023929 | Unknown | | | | | | | | | - | 4.5 | 1.1 | 4.4 |
| 001323 | Protein takeout | | | | | | | | Odorant binding activity | - | - | 8.4 | - |
| 003358 | Unknown | | | | | | | | | - | 3.8 | - | 4.3 |
| 012737 | Nose resistant to fluoxetine protein 6 | | | | | | | | | 1.6 | 4.2 | 1.6 | 3.9 |
| 012164 | SPZ6: spaetzle-like cytokine | | | | | | | | | - | 3.2 | 0.5 | 4.8 |
| 027145 | Unknown (Serpin) | | | | | | | | | - | 4.2 | 0.7 | 3.6 |
| 012642 | Unknown (Chitin binding) | | | | | | | | | - | 3.4 | 0.3 | 4.3 |
| 001498 | Unknown | | | | | | | | | - | 3.4 | 0.5 | 4.1 |
| 025451 | Angiopoietin-related protein 2 | | | | | | | | | - | - | - | 7.0 |
| 015559 | Zinc carboxypeptidase | | | | | | | | Hydrolytic reaction | - | 2.5 | - | 4.5 |
| 025440 | lncRNA | | | | | | | | | - | - | 3.7 | 2.9 |
| 024040 | ATP synthase subunit b, mitochondrial-like | | | | | | | | Proton-transporting ATP synthase complex | - | 6.4 | - | - |
| 002961 | Osiris, putative | | | | | | | | | - | - | 0.9 | 6.3 |
| 003242 | Pupal cuticle protein | | | | | | | | | 6.3 | - | - | -1.4 |
| 003351 | Unknown | | | | | | | | | - | - | 1.0 | 6.0 |
| 005024 | Unknown | | | | | | | | | - | - | 1.1 | 4.6 |
| 010631 | Unknown (Orthologue to Osiris) | | | | | | | | | - | - | - | 5.4 |
| 010628 | Unknown (Orthologue to Osiris) | | | | | | | | | - | - | - | 5.1 |
| 027763 | Lactoylglutathione lyase | | | | | | | | | - | - | - | 4.9 |
| 025469 | Preli-like protein | | | | | | | | | - | - | 4.9 | - |
| 023249 | Unknown | | | | | | | | | - | - | - | 4.9 |
| 004030 | Unknown | | | | | | | | | - | - | - | 4.8 |
| 003337 | Unknown | | | | | | | | | -1.1 | 3.4 | - | 4.7 |
| 014766 | Nose resistant to fluoxetine protein 6 | | | | | | | | | - | - | 4.6 | - |
| 025898 | MAGE-like protein 2 | | | | | | | | | - | - | - | 4.5 |
| 020544 | Pseudogene | | | | | | | | | 4.3 | - | 2.9 | - |
| 001294 | Unknown (Haemolymph_juvenile_hormone-bd) | | | | | | | | | - | - | 3.7 | -1.7 |
| 013031 | APG18A: autophagy related gene | | | | | | | | | - | - | 5.2 | -4.1 |

For each gene, Vectorbase accession, Vectorbase description, predicted function and expression values in the different samples are indicated. Different colour codes indicate genes with a shared function. Significant expression values ($padj < 0.001$) are coloured with a blue/red colour code corresponding to down/up-regulation. "-" means absence from the DESeq2 output of genes with $padj < 0.1$, meaning that transcripts are either not detected or not differentially regulated at this 0.1 threshold.
