## Supplementary material for "Production of germ-free mosquitoes via transient colonisation allows stage-specific investigation of host-microbiota interactions": Table S5

**Table S5**. List of the most down-regulated genes in germ-free larvae compared to colonised larvae.

| AAEL | Description | Amino acid storage | Hydrolytic reaction | Lipid metabolism | Cuticle constituent | Immunity | Function | germ-free vs *E. coli* | germ-free vs *E. coli* | germ-free vs *E. coli* | germ-free vs *E. coli* |
|---|---|---|---|---|---|---|---|---|---|---|---|
| | | | | | | | | GUT_12h | GUT_20h | WL_12h | WL_20h |
| 008045 | Hexamerin 2 beta | ■ | | | | | | - | -6.7 | -5.3 | -7.0 |
| 011169 | Hexamerin 2 beta | ■ | | | | | | -3.6 | -4.2 | -2.5 | -4.2 |
| 011520 | Sucrose transport protein | | | | | | Sucrose transport | -4.5 | -3.8 | -3.1 | -2.9 |
| 008817 | Hexamerin 2 beta | ■ | | | | | | - | -4.0 | -3.1 | -6.0 |
| 018240 | Unknown (Fuseless region) | | | | | | | -3.3 | -3.4 | -3.3 | -3.0 |
| 008313 | Unknown | | | | | | | -2.9 | -3.1 | -4.5 | -4.5 |
| 013757 | Hexamerin 2 beta | ■ | | | | | | - | -5.4 | -2.2 | -4.5 |
| 011661 | Unknown (Cilia structure/activity) | | | | | | | -3.9 | -4.1 | -1.9 | -3.2 |
| 023132 | Unknown (Lipase activity) | | ■ | | | | | -2.7 | -4.0 | -2.5 | -4.5 |
| 008598 | Unknown (Vitellogenin domain) | | | ■ | | | | - | -2.7 | -2.0 | -4.7 |
| 013981 | Hexamerin 2 beta | ■ | | | | | | - | -4.5 | -1.7 | -4.4 |
| 004341 | Carboxy/choline esterase Alpha Esterase | | ■ | | | | | - | - | - | -8.9 |
| 014886 | 4-aminobutyrate aminotransferase | | | | | | | -3.2 | -4.1 | -0.4 | -1.0 |
| 011968 | Unknown | | | | | | | -6.8 | - | - | - |
| 011349 | Serine protease | | ■ | | | | | -1.8 | -3.5 | -0.7 | -1.4 |
| 022253 | Pseudogene | | | | | | | -4.0 | - | -2.5 | - |
| 006824 | Cytochrome P450 | | | | | | Oxidoreductase activity | - | - | - | -6.4 |
| 012311 | Vitellogenin | | | ■ | | | | - | -3.8 | - | -2.3 |
| 002237 | Fatty acid synthase | | | ■ | | | | - | - | - | -6.1 |
| 004343 | OBP19: odorant bp | | | | | | | - | -1.7 | -2.7 | -3.3 |
| 014994 | Unknown (Cuticular protein) | | | | ■ | | Cuticle constituent | -6.7 | - | - | 1.0 |
| 024321 | SWI/SNF complex subunit SMARCC2-like | | | | | | DNA binding | - | -5.5 | - | - |
| 011407 | CTL20: C-Type Lectin | | | | | ■ | | -2.2 | -1.8 | -3.1 | -1.9 |
| 003890 | Cytochrome P450 | | | | | | | - | - | -3.2 | -1.9 |
| 017976 | HSP70Bb: heat shock protein | | | | | | | -3.6 | - | -1.5 | - |
| 024362 | Tetra-peptide repeat homeobox protein 1-like | | | | | | DNA binding | -6.4 | - | -1.6 | 1.6 |
| 013983 | Hexamerin 2 beta | ■ | | | | | | - | -4.5 | -2.2 | -4.7 |
| 011944 | CCEAE3O: carboxy/choline esterase Alpha Esterase | | ■ | | | | | - | - | - | -4.6 |
| 025320 | Cyclin-dependent kinase inhibitor 1C-like | | | | | | Cell cycle regulator | -4.5 | 1.3 | - | 0.8 |
| 000689 | Steroid dehydrogenase | | | | | | | - | - | - | -4.2 |
| 013031 | Unknown (Cuticular protein) | | | | ■ | | | - | - | 5.2 | -4.1 |
| 028198 | Carboxylic ester hydrolase (Fragment) | | ■ | | | | | -4.1 | - | -3.4 | - |
| 009524 | MAL1: alpha-amylase | | ■ | | | | | - | -4.1 | - | -2.4 |
| 021670 | Unknown (Odorant binding protein) | | | | | | | - | - | -3.1 | -4.1 |
| 022059 | Pseudogene | | | | | | | -3.7 | - | -2.0 | - |
| 010134 | Pupal cuticle protein, putative | | | | ■ | | | - | -3.7 | - | - |
| 013152 | Leucine-rich repeat extensin-like protein 3 | | | | | | | -3.5 | - | - | 4.0 |
| 026388 | Cuticle protein 38-like | | | | ■ | | | - | - | -3.4 | - |
| 027210 | Nuclear pore complex protein DDB_G0274915-like | | | | | | Transport into nucleus | -4.9 | - | -0.8 | 2.4 |
| 027233 | Unknown | | | | | | | - | - | -4.3 | 2.7 |
| 000914 | Cuticle protein | | | | ■ | | | - | - | -5.1 | -1.2 |
| 004953 | Elongation of very long chain fatty acids protein | | | ■ | | | | - | 1.5 | -2.8 | 1.9 |
| 022796 | Unknown | | | | | | | - | - | -2.9 | 3.4 |

For each gene, Vectorbase accession, Vectorbase description, predicted function and expression values in the different samples are indicated. Different colour codes indicate genes with a shared function. Significant expression values (padj < 0.001) are coloured with a blue/red colour code corresponding to down/up-regulation. "-" means absence from the DESeq2 output of genes with padj < 0.1, meaning that transcripts are either not detected or not differentially regulated at this 0.1 threshold.
